# Microprotein Regulates G-quadruplex Driven RNA Aggregation

**DOI:** 10.64898/2026.04.10.717804

**Authors:** Bikash R. Sahoo, Janakraj Bhattrai, Arnav Sharma, Ursula Jakob, James CA Bardwell

## Abstract

Repeat expansions of the hexanucleotide GGGGCC in *C9orf72* form aberrant phase transitions that have been linked to Amyotrophic Lateral Sclerosis and Frontotemporal Dementia. RNA structures such as G-quadruplexes and hairpins play important roles in these processes. Here, we show that the human microprotein ZNF706 acts as a modulator of G-quadruplex formation and RNA phase behavior. ZNF706 antagonizes pathological gel-solid transitions by melting hexanucleotide repeat G-quadruplex structures converting gel-like aggregates into more dynamic condensates. Loss of ZNF706 enhances the cellular production clearance of hexanucleotide repeat-mediated dipeptide repeat proteins, while overexpression suppresses their production and promotes clearance. Mechanistically, ZNF706 influences hexanucleotide repeat condensate fluidity and viscoelasticity. We find ZNF706 acts as an RNA chaperone that remodels repeat RNA structures and solubilizes RNA aggregates. This activity represents one mechanism whereby cells can regulate G-quadruplex driven phase transitions linked to neurodegenerative diseases.

## Introduction

The expansion of the hexanucleotide repeat GGGGCC within the non-coding region of the *C9orf72* gene is closely linked with two devastating neurological disorders, Amyotrophic Lateral Sclerosis (ALS) and Frontotemporal Dementia (FTD)^1,2^. This expanded repeat may exert its toxicity through various gain-of-function mechanisms^3^; RAN translation of these repeat RNAs, for instance, can produce toxic dipeptide repeats ^4^. Another recently proposed mechanism involves the formation of stable, self-associating toxic RNA foci composed of the GGGGCC repeats^5,6^. A combination of these two models has also been suggested whereby dipeptide repeats behave like polyamines that promote RNA-RNA interactions and ribonucleoprotein coalescence^7^. In this combined model, expanded-repeat RNAs and their products nucleate condensate formation and enhance their maturation^5,8^. The folding of GGGGCC repeats into highly stable G-quadruplex structures has been implicated in ALS pathology^9–11^. These non-canonical nucleic acid structures are formed by the stacking of planar G-quartets that are stabilized by Hoogsteen hydrogen bonds. GGGGCC repeats can also adopt alternative conformations such as hairpins^12^ which can exist in equilibrium with G-quadruplexes ^13^. The dynamic equilibrium between these secondary structures and their distinct roles in driving cellular dysfunction in neurodegenerative diseases remain poorly understood.

Recent evidence suggests that the structural stability and specific topology of the G-quadruplex RNA within GGGGCC repeats may influence RNA toxicity in *C9orf72* repeat expansion disorders^14^. When transfected into cells, *C9orf72* repeat RNAs can form gel-like condensates in a manner that depends on both their secondary structure and number of repeat units^5,15^.

Several RNA-binding proteins, including FUS^16^ and Zfp106^17^, have been identified as chaperones capable of suppressing RAN translation by modulating the G-quadruplex (G4) structures of GGGGCC repeat RNAs^18^. G-quadruplex containing repeat RNAs can recruit RNA-binding proteins and alter condensate material properties by seeding ribonucleoprotein aggregation^5,19^. Conversely, ribosome binding proteins and helicases that recognize G-quadruplexes may remodel these structures, influencing the translation of RNAs containing G-quadruplexes and broader condensate behavior^20^. In the event that these RNA and proteins are not in appropriate balance, RNAs may coalesce pathologically into nuclear foci^9^. Recent studies have begun to illuminate how G-quadruplex containing RNAs, and ribosome binding proteins co-operate in the formation, modulation, and dissolution of ribonucleoprotein condensates^21^. The molecular strategies by which RNA-binding proteins recognize and suppress GGGGCC repeat RNA aggregation remain to be fully elucidated.

Sequence-specific RNA gelation parallels the way in which protein aggregation contributes to neurodegenerative disease^22^. In both cases, multivalent binding enables phase transitions into higher-order assemblies such as gels or fibrils that disrupt normal cellular functions^5,23^. For RNAs, multivalency can arise from structured motifs (like G-quadruplexes) or from repeat expansions that enable promiscuous RNA-RNA interactions^24,25^. Several factors influence RNA condensation, including ionic strength, RNA secondary structure, and interactions with RNA-binding proteins. Monovalent ribosome-binding proteins and actively translating ribosomes may oppose condensation by competing with RNA-RNA base pairing or by sterically hindering RNA self-association^26^.□Moreover, multivalent RNA-binding proteins or structured repeat RNAs can drive networked assemblies by bridging or scaffolding multiple RNA molecules, promoting gel-like granule formation. Proteins such as nucleolin, the helicases DHX36/Pif1^27,28^, FUS^16^, hnRNP H1^29^, TDP-43^30^, TRF2^31^ and SERF2^32^ have all been shown to interact with RNA G-quadruplexes in vitro to regulate their folding. G-quadruplex elements may link RNA secondary structure to phase behavior by seeding multivalent interactions, recruiting ribosome binding proteins, or both.

Above a critical repeat length, expanded-repeat RNAs undergo multivalent base-pairing that results in a liquid-to-gel or liquid-to-solid phase transition. This phenomenon can also drive RNA foci formation in cells^5,11^. G-quadruplex RNA foci are observed abundantly in fibroblasts derived from patients with C9 ALS^33^.

A handful of RNA binding proteins have been implicated in counteracting GGGGCC-mediated pathology by targeting G-quadruplexes in RNA. For instance, FUS, which has been linked to ALS/FTD, binds directly to GGGGCC RNAs and remodels their G-quadruplex conformations, thereby suppressing RAN translation^10,16,17,34^. Zfp106, which is essential for motor neuron survival) acts similarly, binding GGGGCC RNAs and inducing conformational change. Overexpression of Zfp106 causes a shift in GGGGCC G-quadruplex structure and markedly reduces both RNA foci and dipeptide repeat protein production in cell models^17^. These findings suggest that G-quadruplex binding proteins can antagonize repeat-RNA condensation and non-canonical translation.

Here, we focus on human ZNF706, a microprotein homologous to the SERF family of proteins^35^. We previously found that SERF proteins specifically bind to G-quadruplexes^32,35^. To extend the paradigm of a microprotein-RNA G-quadruplex interaction into a disease-relevant area, we investigated the interaction of ZNF706 with GGGGCC repeats. We show here that ZNF706 modulates translation and condensate dynamics in repeat-expansions by recognizing G-quadruplexes formed from GGGGCC RNA repeats, promoting solubilization of repeat RNA aggregates, in turn suppressing RAN translation and reducing dipeptide repeat protein accumulation.

## Results

### ZNF706 shows affinity for long repeat RNA G-quadruplexes

Human ZNF706 is a 76 residue microprotein composed of an N-terminal intrinsically disordered SERF homologous domain (Figure 1) fused to a C-terminal C2H2 zinc-finger (ZNF)^35^. We earlier showed that ZNF706 specifically binds G-quadruplex structures, including some formed by pathologically relevant RNA repeats^35^. As such, we elected to test whether ZNF706 recognizes the pathogenic GGGGCC repeats associated with *C9orf72*-linked neurodegeneration; GGGGCC is known to form RNA G-quadruplex structures (Figure 1)^11^. We found ZNF706 binds a short 4-repeat GGGGCC, with a binding dissociation constant of 1.7 ± 0.2 µM. ZNF706 shows a similar (0.6 ± 0.2 µM) binding affinity for a 4-repeat telomeric repeat RNA (UUAGGG), another well-characterized RNA G-quadruplex^36^ (Figure S1a, Table 1). As the quantity of the GGGGCC repeat is increased, so does its binding affinity, plateauing out for long 32-repeat or disease relevant 69-repeat GGGGCC transcripts^11^ with binding dissociation constants in the low nanomolar (∼30 nM) range (Figure S1b, c and Table 1). Binding was negligible when these RNAs were mutated (UUACCG or CCCCGG) to disrupt G-quadruplex formation (Figure S1a). ZNF706 also binds to DNA oligonucleotides of the same GGGGCC repeat sequences, albeit with a similar or slightly reduced affinity (Figure S1d).

**Figure 1.**
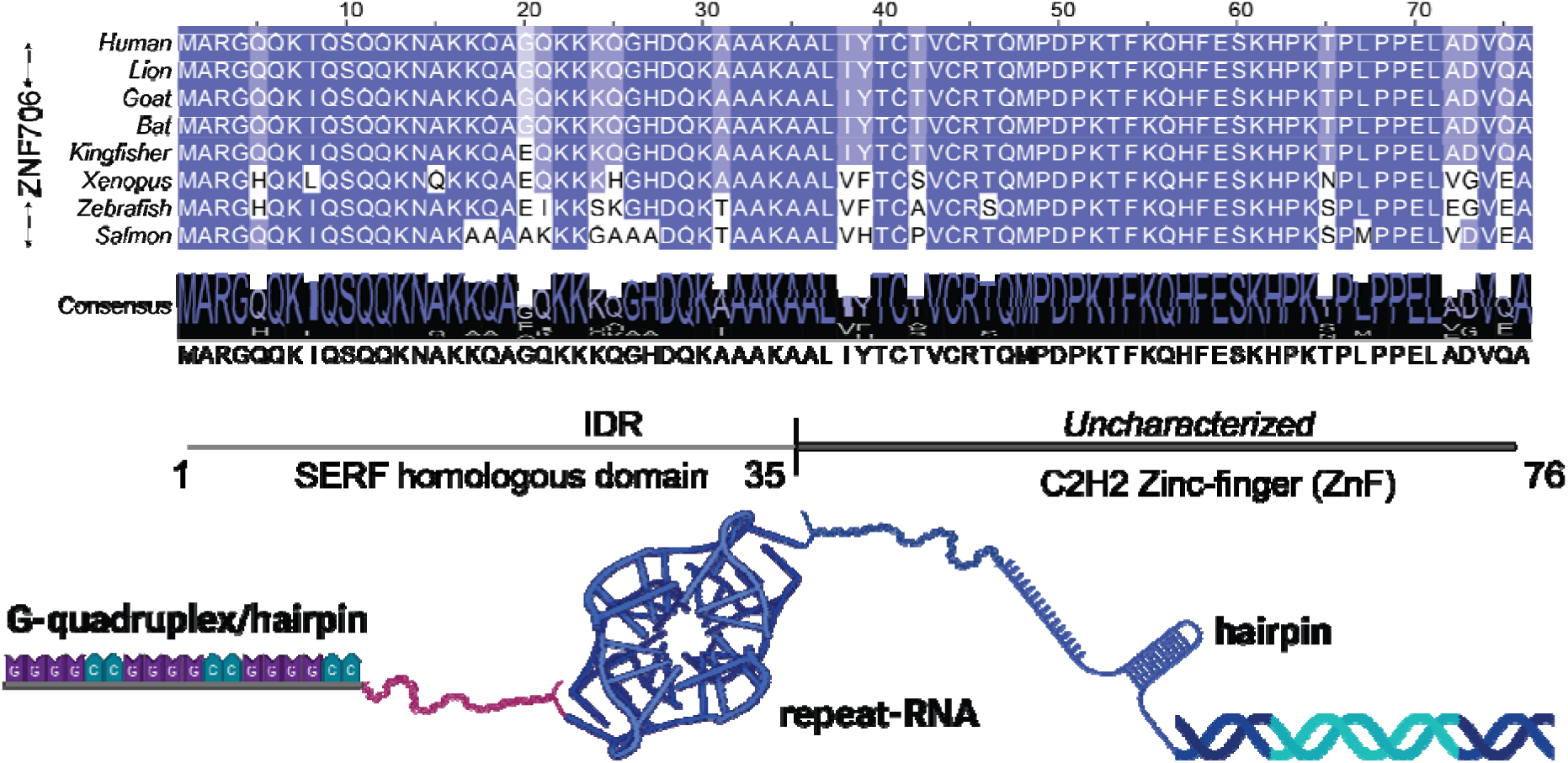
ZNF706 and GGGGCC repeat RNA architecture. Multiple sequence alignment of ZNF706 across representative vertebrate species including humans shows strong conservation across the species. Domain organization of human ZNF706 is illustrated below, highlighting its intrinsically disordered N-terminal region (IDR) followed by a C-terminal C2H2 zinc-finger (ZnF) domain. A schematic representation of GGGGCC repeat RNA is shown forming higher-order structures, including G-quadruplexes and hairpins. This schematic summarizes the conceptual framework of the study, in which ZNF706 interacts with structured repeat RNA derived from expanded *C9orf72* GGGGCC sequences. This image is created with BioRender.com.

**Table 1.**
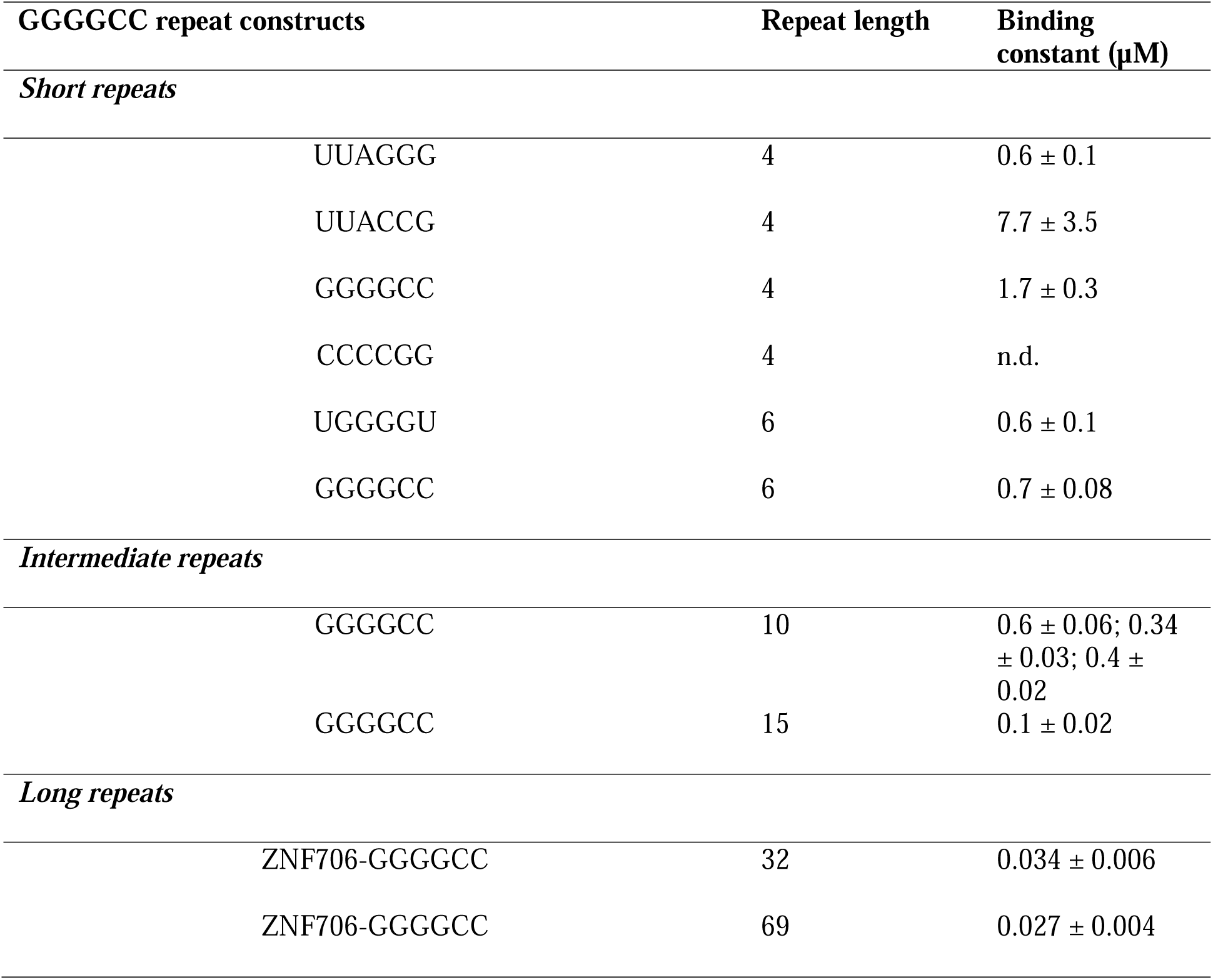
ZNF706 binds tighter to disease-relevant long GGGGCC RNA repeats.

| GGGGCC repeat constructs | Repeat length | Binding constant ($\mu\text{M}$) |
| --- | --- | --- |
| *Short repeats* |  |  |
| UUAGGG | 4 | $0.6 \pm 0.1$ |
| UUACCG | 4 | $7.7 \pm 3.5$ |
| GGGGCC | 4 | $1.7 \pm 0.3$ |
| CCCCGG | 4 | n.d. |
| UGGGGU | 6 | $0.6 \pm 0.1$ |
| GGGGCC | 6 | $0.7 \pm 0.08$ |
| *Intermediate repeats* |  |  |
| GGGGCC | 10 | $0.6 \pm 0.06$; $0.34 \pm 0.03$; $0.4 \pm 0.02$ |
| GGGGCC | 15 | $0.1 \pm 0.02$ |
| *Long repeats* |  |  |
| ZNF706-GGGGCC | 32 | $0.034 \pm 0.006$ |
| ZNF706-GGGGCC | 69 | $0.027 \pm 0.004$ |

Across three different GGGGCC-repeat G-quadruplexes, ZNF706 shows a repeat length-dependent increase in binding affinity (Figure S1e) similar to what was observed for RNA repeats. ZNF706 appears relatively agnostic towards an RNA or DNA backbone, instead recognizing the G-quadruplex architecture itself, evidenced by the fact that a very similar binding affinity of ∼700 nM was measured for a 6-repeat UGGGGU and GGGGCC G-quadruplex sequences (Figure S1e, f).

ZNF706-RNA complexes can also be detected using a native gel electrophoretic mobility shift assay for both GGGGCC and UUACCG G-quadruplexes. (Figure S1g, h). These binding studies characterize ZNF706 as an RNA G-quadruplex binding microprotein that shows a similar binding affinity to G-quadruplex sequences as are found for other much larger and biophysically less amenable RNA G-quadruplex binding proteins such as FUS or TDP-43^37–39^.

### ZNF706 increases the fluidity of GGGGCC RNA foci in cells

As ZNF706 binds to GGGGCC repeat RNA G-quadruplexes in vitro (Table 1), we decided to investigate its role in cellular models in *C9orf72* pathology. In *C9orf72*-associated ALS and FTD, the GGGGCC repeat expansion has been linked to toxicity via both repeat RNA aggregation and translation of dipeptide repeat protein. Multivalent G-quadruplex associations allow these repeat RNAs to undergo length-dependent phase transitions, and it has been hypothesized that RNA condensation may represent an early, disease-relevant event ^40,41^.

ZNF706’s ability to bind the RNA G-quadruplexes formed by GGGGCC repeats represents one possible way for ZNF706 to affect GGGGCC mediated aggregation. RNA G-quadruplexes can form cellular foci whose transition to a gel-like state leads to the sequestration of key RNA binding proteins, possibly contributing to the toxicity of GGGGCC repeats^40^. We thus explored if ZNF706 could influence the localization and material properties of cellular foci formed by GGGGCC repeats. We used a previously described monoclonal U2OS cell line stably expressing a MS2-YFP fusion that can report on the solubility of MS2-tagged repeat RNA transcripts^5^. These cells were transfected with a plasmid encoding 34-repeat GGGGCC, resulting in the formation of distinct, punctate nuclear RNA foci (Figure 2a, b) as previously reported^5^. ZNF706 overexpression showed diffuse nuclear localization and did not prevent the formation of distinct, punctate GGGGCC RNA foci (Figure 2c, d), indicating that elevated ZNF706 levels are insufficient to block repeat RNA condensation. The persistence of morphologically similar nuclear foci prompted us to investigate whether ZNF706 alters the material properties of GGGGCC assemblies rather than their formation.

**Figure 2.**
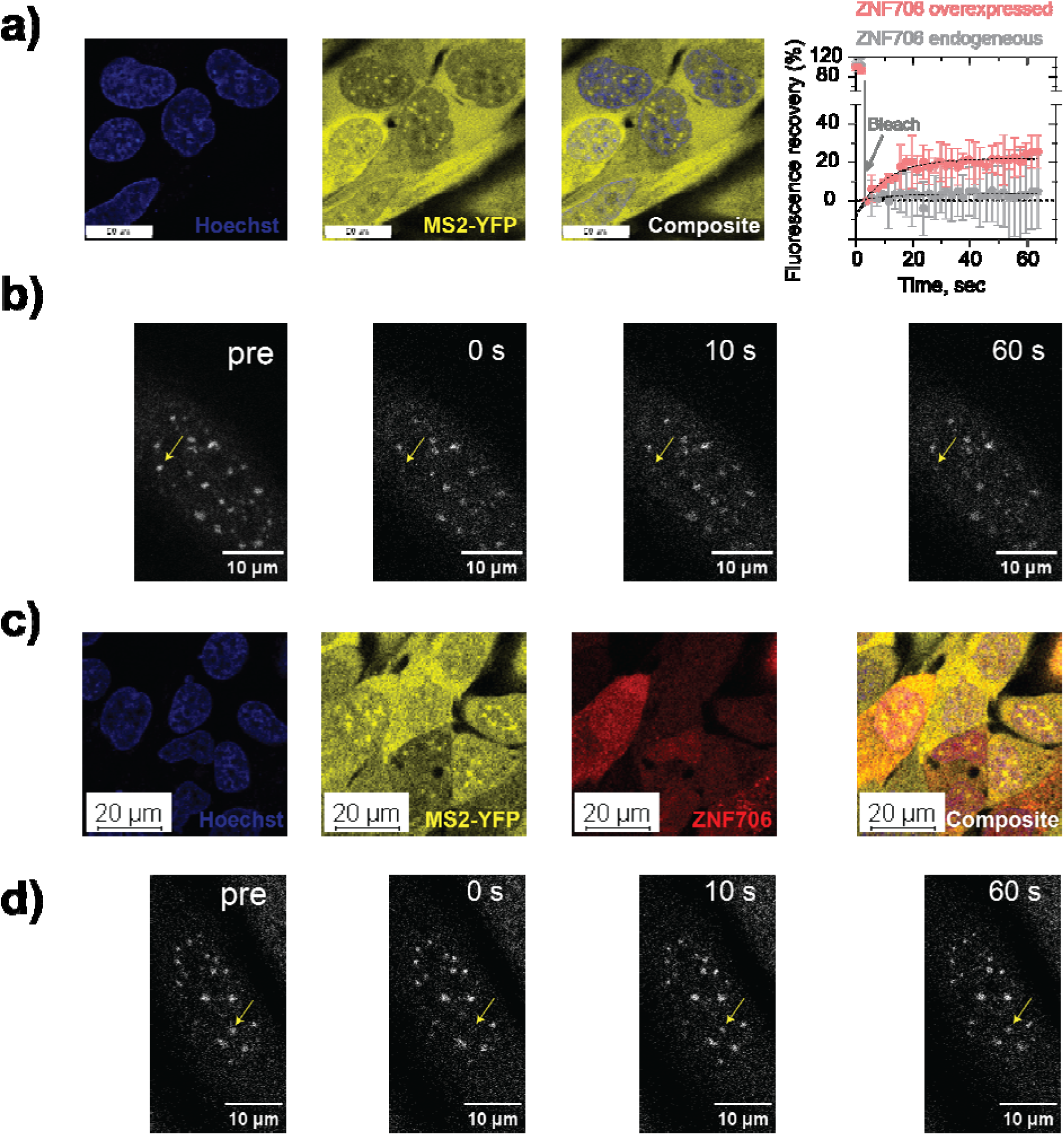
ZNF706 regulates GGGGCC RNA foci dynamics. (a,c) Live-cell imaging of monoclonal U2OS cells that stably express MS2-YFP protein (yellow) show discrete RNA foci within in nucleus (blue, Hoechst stain) formed by MS2-tagged 34×GGGGCC RNA transcripts in cells expressing endogenous ZNF706 (a), and cells overexpressing (c) ZNF706-mCherry (red). Scale bar, 20 µm. The panel on the top right shows the quantitative FRAP recovery plots (mean ± s.e.m., n = 3) corresponding to panels (a) and (c) for cells transfected without or with ZNF706-mCherry. (b,d) FRAP measurement time series showing post-bleach recovery of GGGGCC RNA foci in the absence (b) or presence (d) of ZNF706-mCherry expressing cells.

To assess if the molecular exchange dynamics and material properties of GGGGCC foci were changed following ZNF706 overexpression, live-cell Fluorescence Recovery After Photobleaching (FRAP) experiments were performed. GGGGCC repeat RNA-foci formed in control cells expressing endogenous levels of ZNF706 exhibited very little fluorescence recovery over the 60-second measurement period showing that these RNA foci exist in a highly static, aggregate-like state (Figure 2a, gray curve), possibly reflecting a matured state^40^. Overexpression of ZNF706-mCherry resulted in partial fluorescence recovery with a half-time of ∼7 seconds (Figure 2a, red curve), demonstrating enhanced molecular mobility and material exchange within GGGGCC foci upon ZNF706 overexpression.

### ZNF706 cellular level regulates GGGGCC repeat-mediated dipeptide repeat protein production

Given ZNF706’s high-affinity binding to GGGGCC repeat RNA, we asked whether these interactions might also influence the degradation or translation of dipeptide repeat protein. We first examined the impact of ZNF706 depletion on the steady state levels of poly-glycine-alanine (Poly-GA) dipeptide repeat protein, which are generated via GGGGCC repeat-associated non-AUG (RAN) translation a process which is dependent on the formation of G-quadruplex structures within the RNA transcript^17,42^. We expressed poly-GA dipeptide repeat protein of varying lengths (GA3, GA35, and GA70) in HEK293 cells. We found that the levels of these dipeptide repeat proteins were elevated (∼2-3 fold) following depletion of ZNF706 (Figures 3a and S2a).

**Figure 3.**
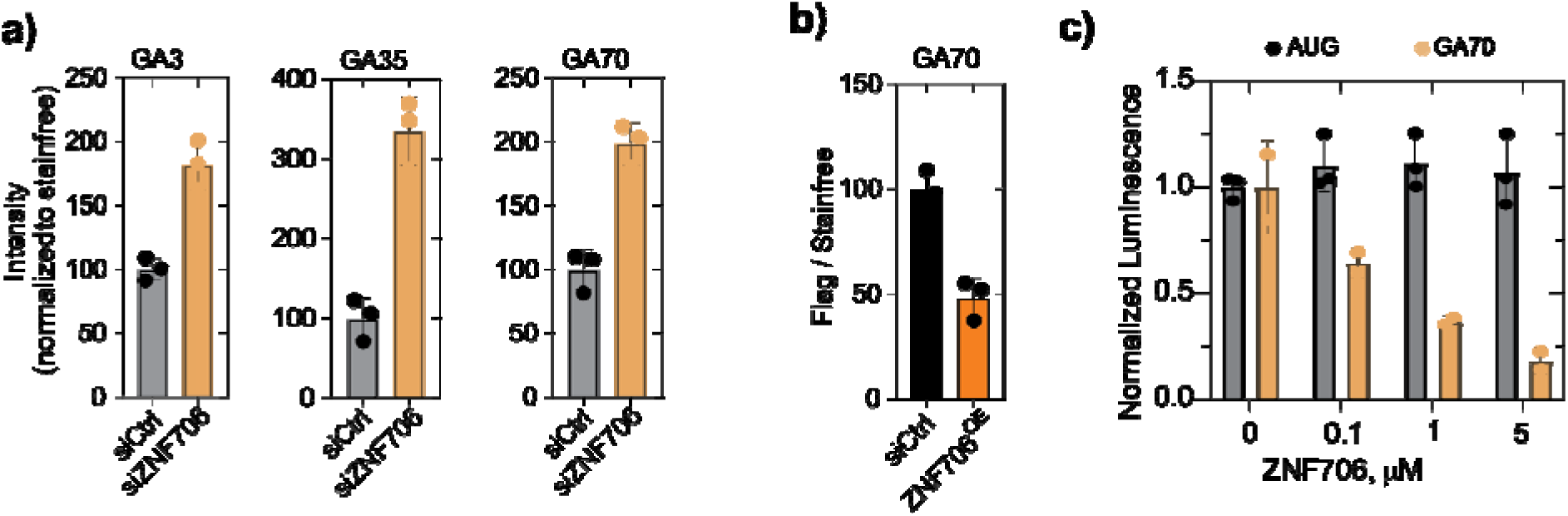
ZNF706 suppresses *C9orf72* dipeptide repeat protein accumulation. **(a)** ZNF706 depletion significantly elevates dipeptide repeat protein levels relative to control. Quantification of GA3, GA35 and GA70 immunoblot signals, normalized to total protein as measured by stain-free SDS-PAGE are shown in Figure S2a. Representative immunoblots showing the levels of poly-GA dipeptide repeat proteins (GA3, GA35, GA70) in HEK293 cells following a control knockdown (siCtrl) or ZNF706 depletion (siZNF706) and stain-free gel images are also shown in Figure S2a. Data represent independent biological replicates (n=3) with mean ± s.d. **(b)** ZNF706 overexpression reduces steady-state levels of GA70. Quantification of GA70 levels normalized to stain-free signal, (Figure S2b) showing decreased GA abundance in the cells overexpressing ZNF706. Representative immunoblots of GA70-Flag expressed in cells with or without ZNF706 overexpression, probed with anti-Flag or anti-GA antibodies are also shown in Figure S2b. **(c)** In vitro translation assay showing normalized luminescence from an AUG-initiated control reporter or a GA70 repeat reporter in the presence of increasing concentrations of purified ZNF706. ZNF706 selectively suppresses translation of the repeat-derived GA70 reporter while minimally affecting canonical AUG-initiated translation.

To complement these knockdown assays, we studied whether overexpression of ZNF706 is sufficient to reduce repeat protein accumulation. We transduced HEK293 cells with lentiviral vector expressing ZNF706 following transfection with GA70-encoding plasmid and compared dipeptide repeat protein levels. ZNF706 overexpression decreased poly-GA levels by about ∼50-60% (Figures 3b and S2b). These changes in dipeptide repeat protein levels upon altering ZNF706 levels could stem either from decreased degradation or increased production, or both. To test these possibilities, we first used cycloheximide to inhibit *de novo* protein synthesis. ZNF706 depletion led to increased polyGA levels at the start of the cycloheximide chase, which remained elevated throughout the time course (Figure S2c,d). Although control cells exhibited a reduction in polyGA signal over time, the persistence of higher levels in ZNF706-depleted cells suggests either increased initial production or altered turnover. However, the results of these chase experiments are not clear enough to make definitive conclusions. In an in vitro translation assay, ZNF706 showed a dose-dependent suppression of GGGGCC-mediated translation (Figure 3c). Canonical AUG-initiated translation remained unaffected, indicating that ZNF706 specifically modulates RAN translation in vitro.

In summary, our cellular studies indicate that ZNF706 may counteract repeat-RNA pathology by suppressing repeat-driven translation and thereby limiting dipeptide repeat protein accumulation. Other RNA-binding proteins are also known to influence repeat RNA structure and RAN translation in various ways. The G-quadruplex helicase DHX36 enhances RAN translation by unwinding RNA G-quadruplex structures ^34^. FUS remodels G-quadruplexes to repress RAN translation^16^. FMRP and TDP-43 modulate translation through direct G-quadruplex binding, affecting ribosomal engagement and mRNA stability^30,43^. An attractive hypothesis is that ZNF706’s effects are due to its ability to bind to G-quadruplexes that may modulate repeat RNA conformation and condensate properties, influencing their translational competence.

### ZNF706 condensates with GGGGCC RNAs exhibit repeat-dependent morphology and mechanics

The live cell experiments suggest that ZNF706 may enhance the fluidity of RNA foci. To explore the fluidity issue in more detail, we turned to in vitro experiments. We first studied whether repeat length and G-quadruplex formation in GGGGCC repeats could influence the material properties of ZNF706-RNA condensates. We found that ZNF706 could form dynamic droplets, in crowded conditions, in the presence of total HeLa cell RNA or repeat RNA G-quadruplexes (Figures 4a-e and S3). Droplets formed by mixing ZNF706 with a short 4-repeat GGGGCC showed faster FRAP recovery times (∼30 sec) similar to recovery times observed in total RNA than those formed by mixing ZNF706 with longer 15- or 20-repeats, where recovery half-time extended to ∼200 seconds (Figure 4b, d, f). Thus, increasing repeat length shifts condensate behavior towards slower dynamics. In addition, the 15R droplets frequently contained internal clusters were not observed in 4-repeat condensates (Figure 4e, inset), suggesting that increased structural heterogeneity is present within longer repeat assemblies. In the absence of ZNF706, long repeat RNAs formed visible aggregates under crowded conditions (Figure S3, right panel). These observations are consistent with the idea that longer GGGGCC repeat RNAs may introduce multivalent interactionsClick or tap here to enter text. that promote internal structuring and influence condensate organization and dynamics ^5^.

**Figure 4.**
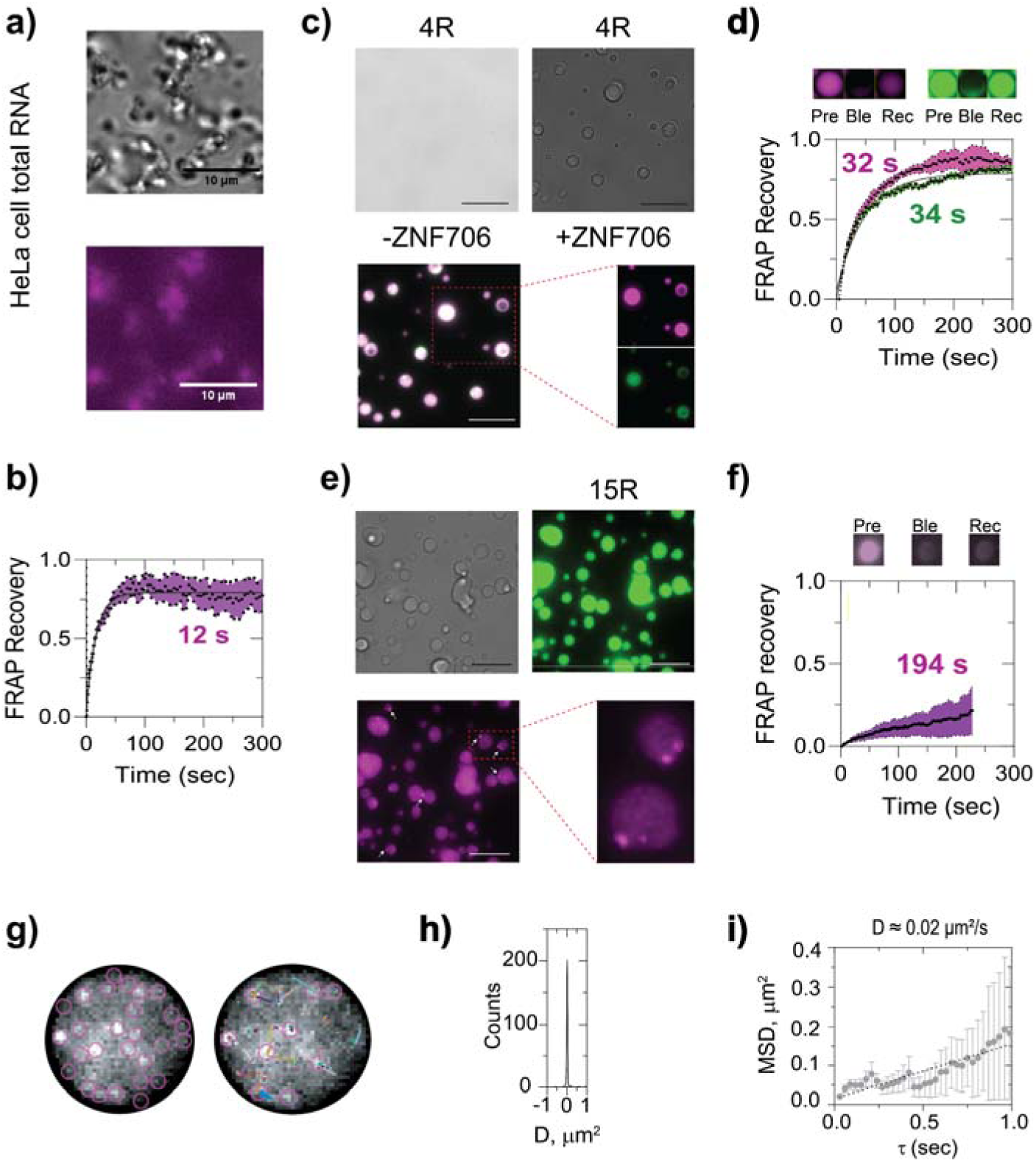
ZNF706 co-condenses with GGGGCC repeats. **(a)** ZNF706 phase-separates with total HeLa RNA above 3.1□μM, forming irregular condensates. **(b)** FRAP analysis shows rapid recovery (t□/□ ≈ 12□s, n = 8), consistent with liquid-like dynamics despite morphological asymmetry. **(c-d)** 4-repeat GGGGCC RNA G-quadruplex show no droplets (left) but forms spherical condensates with ZNF706 (purple) that displays intermediate recovery kinetics (t□/□ ≈ 30□s, n = 8) (d). **(e-f)** ZNF706 mixed with 15× GGGGCC RNA G4 forms droplets that exhibit internal nanodomain patterning (arrows) and slower dynamics (t□/□ ≈ 194 s, n = 8) (f), consistent with emergent mesoscale crosslinking. The sample mixture in (c and e) contained a 1200:200 1 ratio of unlabeled to Cy5 unlabeled protein (Cy5) or FAM labelled RNA. (FAM). **(g)** Representative fluorescence images of a single ZNF706-GGGGCC 15-repeat RNA droplet containing picomolar ZNF706-Cy5 was analyzed to calculate the diffusion of ZNF706 by single-particle tracking in ImageJ. **(h-i)** Plots showing the diffusion constant (h), and the mean square displacement (i), of ZNF706 in the protein-RNA condensate in theformed using a 15-repeat GGGGCC.

To quantify ZNF706’s mobility’s dependence on repeat length, we performed single-particle tracking in 15R and 4R RNA condensates (Figure 4g). The diffusion coefficient (D) for ZNF706 in 15R droplets was a slow 0.02 µm²/s (Figure 4h, i),^32,44,45^ The mobility in 4R condensates was about ∼ 5-fold higher (Figure S4a). These rates are within the range of ∼0.01-0.6 µm²/s that has been found for other proteins present in ribonucleoprotein condensates^32,44,45^ Our data indicate ZNF706 remains dynamic in both short and long repeat RNA condensates, although mobility is substantially reduced in the latter.

We next probed whether these differences in molecular mobility are reflected in the bulk material properties of the condensates. In simple liquid droplets, surface tension drives rapid spherical coalescence^46^. More viscous droplets resist fusion and deform elastically. To test this principle in our system, we measured droplet fusion dynamics using optical tweezers. Fusion times increased with repeat length, with 15R droplets fusing much more slowly than 4R droplets (Figure 5a). This repeat-length dependent increase in fusion time indicates a transition from a simple liquid to a more viscoelastic fluid, where internal polymer interactions resist coalescence.

**Figure 5.**
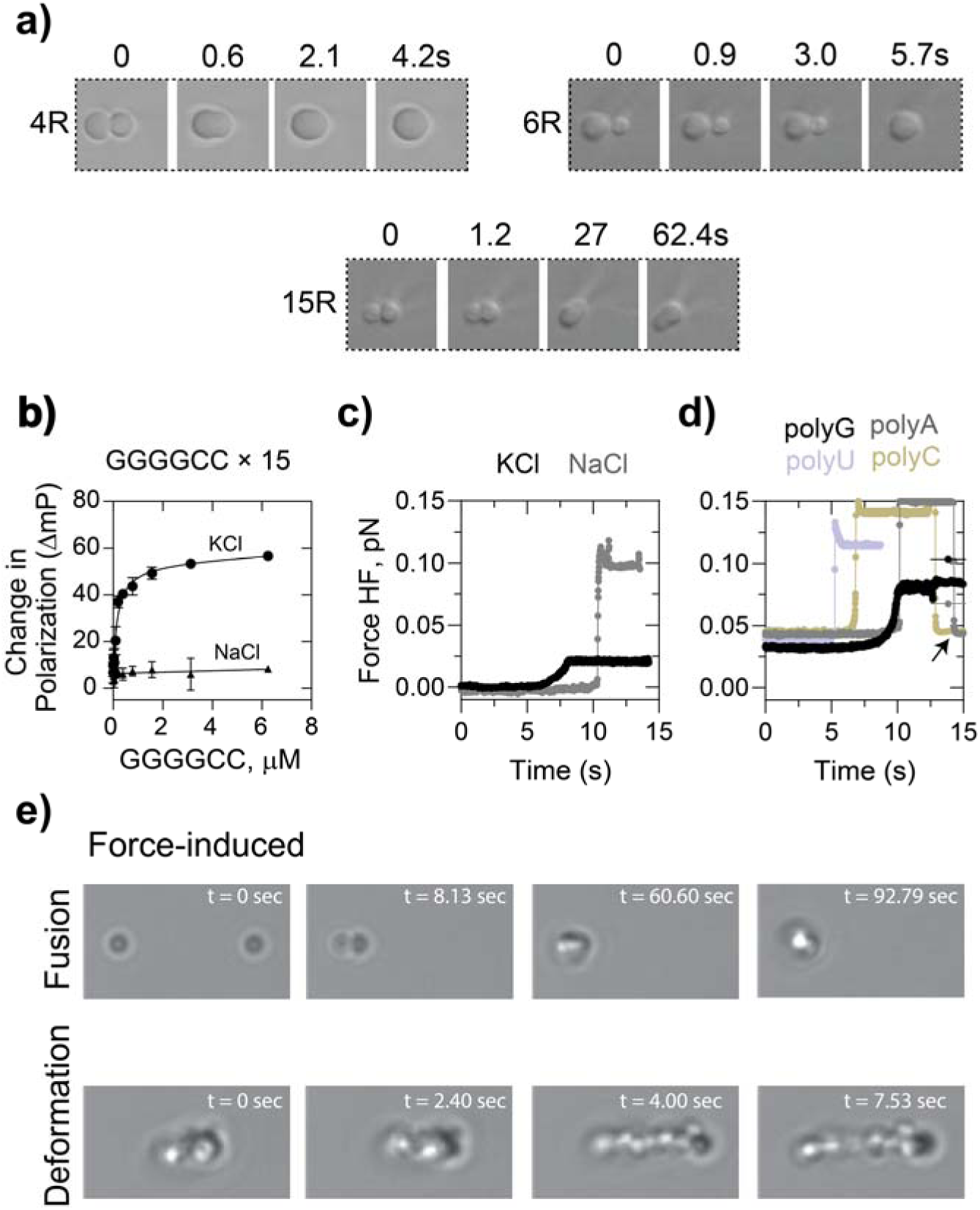
Expanded GGGGCC repeat RNAs form viscoelastic ZNF706-RNA condensates whose mechanical properties depend on repeat length and ionic conditions. **(a)** Time-lapse differential interference contrast (DIC) microscopy showing fusion dynamics of ZNF706-RNA condensates formed using GGGGCC repeats of different lengths: 4R, 6R and 15R. Short repeats undergo rapid droplet fusion within seconds, whereas longer repeats exhibit slower coalescence dynamics, consistent with increased material rigidity. Time points are indicated above each frame. **(b)** Fluorescence polarization measurements of 15× GGGGCC RNA as a function of RNA concentration in the presence of KCl or NaCl. Increasing RNA concentration promotes higher polarization in KCl, indicative of assembly into higher-order structures, whereas NaCl produces minimal change, consistent with ion-dependent structural stabilization. **(c)** Representative force traces from mechanical perturbation experiments showing the response of GGGGCC RNA condensates in buffers containing KCl or NaCl. Condensates formed in KCl exhibit a pronounced force response compared to NaCl in fusion and pull-force deformation, indicating greater mechanical resistance. **(d)** Force versus time profiles of condensates formed by homopolymeric RNAs (polyG, polyA, polyU and polyC). PolyG assemblies display a distinct force response compared to other RNA homopolymers, consistent with the formation of structured assemblies such as G-quadruplex-stabilized networks. **(e)** Time-lapse DIC microscopy illustrating force-induced remodeling of RNA condensates. Upper panels show droplet fusion following mechanical perturbation. Lower panels show deformation and elongation of condensates under sustained force, highlighting the viscoelastic nature of repeat RNA assemblies. Time stamps are indicated for each frame.

### ZNF706 condensate material properties are driven by RNA G-quadruplex folding

We next asked if RNA G-quadruplex structure underlies the increased solidity of long-repeat assemblies. We investigated this by examining how ionic conditions and RNA sequence influence fusion dynamics and droplet deformability. Short GGGGCC repeats form stable G-quadruplex structures, whereas long GGGGCC repeats show equilibrium between G-quadruplex and hairpin structures^13,47^. Additionally, GGGGCC RNAs are favored to fold into G-quadruplex structures in the presence of potassium ions and into hairpins in the presence of sodium^39^ (Figure S4b). Changing the cation of the buffer offers a way to modulate structure while keeping the RNA sequence constant. We initially tested the effect of these cations on 15 and 32-GGGGCC repeats. CD spectroscopy shows that these repeats folded in KCl solution have sharp positive and negative bands at ∼264 and 240 nm, respectively, that were absent in water or NaCl solution correlating to previous observations for short repeats^48^. These significant differences are indicative of KCl solution-mediated G-quadruplex formation (Figure S4b). ZNF706 and 15 repeat GGGGCC co-condensates remained slow-fusing and elastic in KCl (Supporting videos SV1-SV2 and Figure S4c). Substituting KCl with NaCl, which favors hairpin structure and ZNF706 binding, restored rapid fusion and fluid-like behavior (Figure 5b, c), implicating G-quadruplex folding mediating slow condensate fusion.

A similar trend was observed when condensate fusion was studied in 4×GGGGCC versus 4×CCCCGG repeats (Supporting videos SV3-SV4).

Poly(G) RNA folds into G-quadruplexes,^49^ while poly(A), poly(U), or poly(C) RNA do not. Droplets formed from ZNF706 Poly(G) mixtures showed delayed fusion kinetics similar to kinetics found for GGGGCC. ZNF706 poly(A/U/C) RNA all supported rapid condensates fusion (Supporting videos SV5-SV7), providing more evidence that G-quadruplex-encoding sequences likely support slow fusion in the presence of ZNF706 (Figures 5d and S5).

To probe material mechanics, we again applied optical tweezer-based deformation. Upon fusion, 20-repeat GGGGCC-ZNF706 condensates deformed into extended gel-like strands (Figure 5e), consistent with a viscoelastic network (Supporting videos SV8-SV9). In contrast, fused condensates of poly(A/U/C) RNA resisted deformation, remaining spherical, while poly(G)-ZNF706 condensates showed an intermediate gel-like behavior (Figure S5, left panel). GGGGCC repeat RNAs may form multivalent G-quadruplex clusters that can crosslink ZNF706 into a network that locally arrests diffusion and slows fusion. Disrupting G-quadruplex structure appears sufficient to dissolve such condensates^50^.

### ZNF706 remodels GGGGCC RNA aggregates into dynamic condensates

Given ZNF706’s ability to bind long GGGGCC G-quadruplexes (Figure S5, right panel) and form dynamic co-condensates, we asked whether ZNF706 could remodel preformed RNA aggregates that had been made using GGGGCC repeats of differing lengths. We induced aggregation of G-quadruplexes formed from GGGGCC repeats in vitro using 10% PEG8000. As expected, RNAs containing intermediate repeat lengths 15 to 32 repeats formed solid-like aggregates (Figures 6a and S3, right). Addition of ZNF706 at a 5-fold molar excess relative to RNA converted these aggregates into spherical fluid droplets (within 30 minutes) that were fluid as indicated by short FRAP recovery times (Figure 6a). Even in over 40-fold molar excess, however, ZNF706 was incapable of dissolving 69R repeats (Figures 6b and S6). ZNF706 however was able to dissolve aggregates in 8- and 35-repeat RNAs (Figure 6b). Condensates formed by ZNF706 dissolving 35-repeat RNA aggregates show FRAP recovery at ∼52 seconds, while 69-repeat RNA structures showed minimal recovery in the presence of ZNF706 (Figure 6c). This length threshold roughly aligns with the repeat lengths of ∼ 24-100 repeats that have been associated with disease^11^.

**Figure 6.**
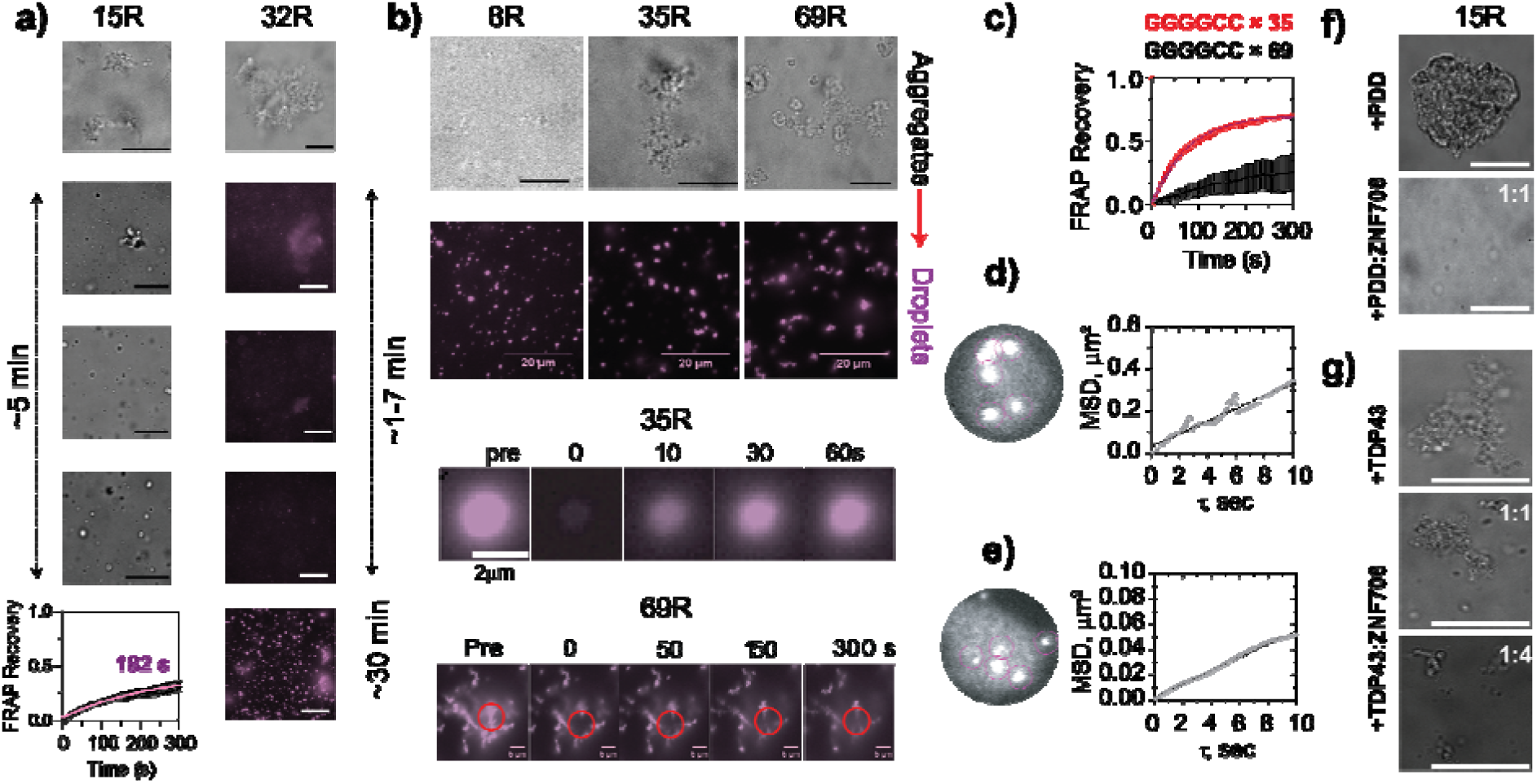
ZNF706 actively remodels GGGGCC RNA aggregates into repeat-length-dependent, dynamic condensates. **(a)** Phase-contrast and fluorescence images of 10 µM 15R and 32R GGGGCC RNA incubated in 10% PEG8000 show large, solid-like aggregates. Upon addition of ZNF706 to a final concentration of (50LJµM,), these assemblies transition into a heterogeneous mixture of punctate droplets and residual aggregates within 5-30LJmin. FRAP analysis of ZNF706-15R condensates (bottom left) reveals partial fluorescence recovery (with a recovery half-time ≈ of ∼182LJs180 s; mean ± s.e.m., nLJ=LJ8), indicating dynamic exchange. Scale bar is 20 µm. **(b)** Representative brightfield and fluorescence images show 8R, 35R, and 69R GGGGCC RNAs (0.5LJµM) in the absence or presence of ZNF706 (5 µM for 8R, and 20LJµM for 35R or 69R). The 8R and 35R RNAs phase separate into uniform, spherical condensates upon ZNF706 addition, while the longer 69R RNA retains remains as large aggregates. Scale bar is 20 µm. Confocal FRAP of ZNF706-35R droplets shows faster recovery (recovery half-time ≈ 52LJs), while 69R aggregates show negligible recovery (bottom panels). **(c)** Quantitative FRAP curve of 35R and 69R RNA condensates formed from pre-aggregated RNA by ZNF706 remodeling. Recovery kinetics (mean ± s.e.m., n = 8) are consistent with the formation of viscoelastic condensates upon ZNF706 addition. **(d-e)** Single-particle tracking of 500LJnm beads embedded in ZNF706 condensates formed with either 4R (d) or 15R (e) RNA GGGGCC RNAs. Mean squared displacement (MSD) curves (mean ± s.d.) demonstrate reduced diffusion and increased viscosity in 15R droplets. **(f-g)** ZNF706 addition induces noticeable disaggregation within 5 minutes of GGGGCC repeat RNA aggregates Disaggregation effect of ZNF706 on GGGGCC repeat RNA aggregates (∼within 5 minutes)that had been formed with the G-quadruplex stabilizing small-molecule pyridostatin (PDD) (f) or a fragment of TDP43

Single particle microrheology, which tracks the motion of embedded biomolecules, revealed that condensate viscosity increased steeply with GGGGCC repeat length. Droplets containing ZNF706 and 4-repeat transcripts of GGGGCC were ∼120 times more viscous than water (Figure 6d), whereas 15-repeat GGGCC-ZNF706 condensates were ∼ 740 times more viscous than water (Figure 6e), a value that surpasses the viscosity previously measured for multivalent peptide-RNA condensates. ZNF706also dissolved 15-repeat RNA aggregates that had been formed in the presence of the G-quadruplex-stabilizing ligand pyridostatin (Figure 6f) and reduced aggregate size when added to TDP-43 RRM1-2/GGGGCC repeat assemblies (Figure 6g). Our results show that ZNF706 can remodel aggregates in a diverse set of circumstances.

### Mechanistic understanding of ZNF706 binding to repeat RNA

We determined the ZNF706 and 4-repeat GGGGCC RNA interaction interface by mapping the folded G-quadruplex binding interface onto our previously solved structure of ZNF706^35^ (Figure 7a). Per-residue amide chemical shift perturbations (CSPs) upon RNA binding reflect changes in the residue’s local environment, with larger shifts suggestive of closer contact (Figure S7, left). Chemical shift perturbations mapped almost exclusively to the disordered N-terminal region of ZNF706 (Figures 7b and S7), indicating direct engagement of the G-quadruplex by the disordered N-terminus, with little or no contribution from the C-terminal zinc finger, consistent with our prior studies^35^.

**Figure 7.**
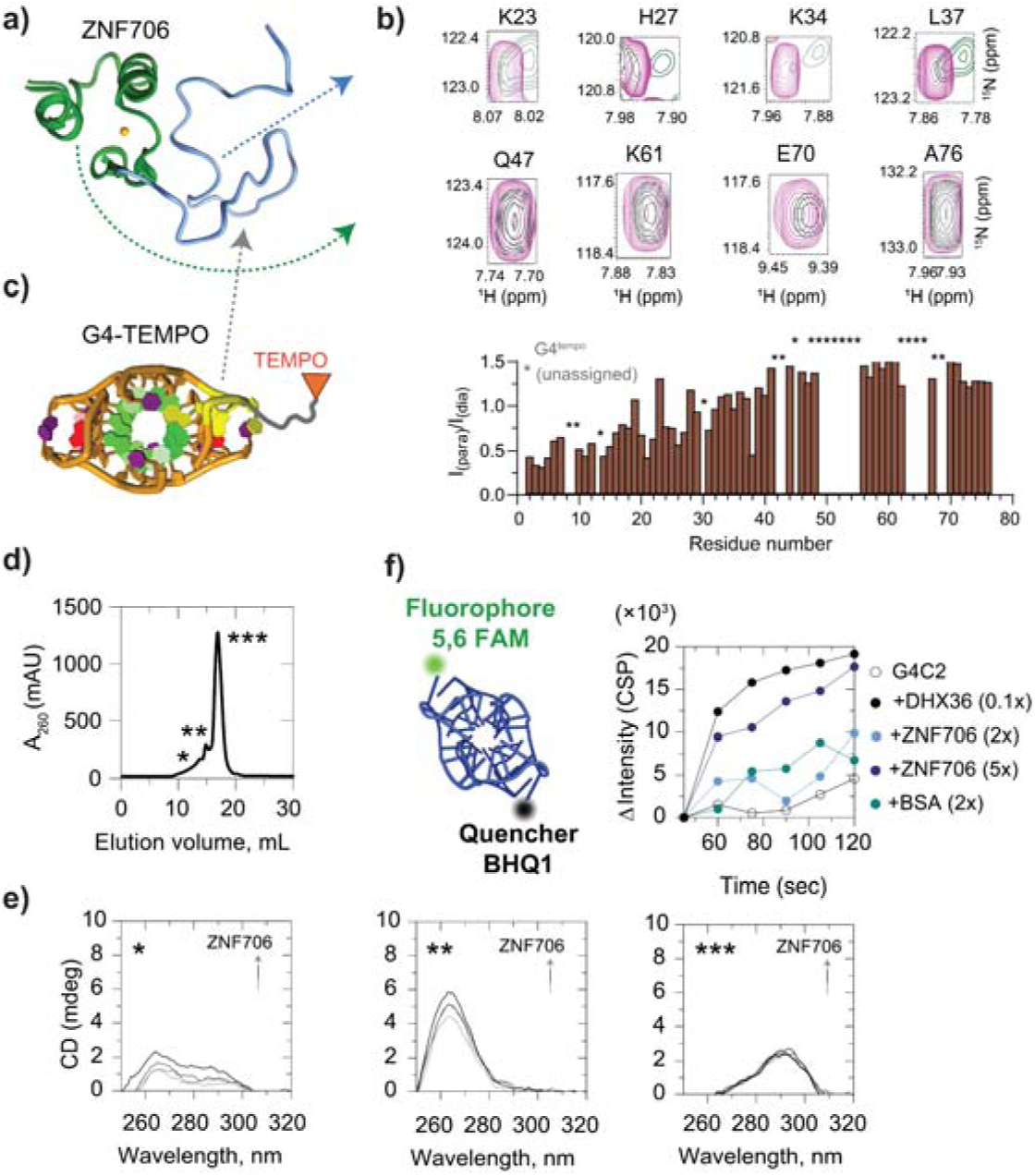
ZNF706 binds and destabilizes parallel GGGGCC G-quadruplexes via its disordered N-terminal SERF-like domain. **(a)** (a) Structure of ZNF706 showing a disordered N-terminal region homologous to SERF proteins and a structured C-terminal C2H2 zinc finger. **(b)** Selected 1H-15N HSQC spectral overlays showing chemical shift perturbations (CSPs) in individual residues of ZNF706 (100 µM) upon titration with GGGGCC repeat RNA G-quadruplex (50 µM), highlighting interactions occur primarily in with the N-terminal domain (full overlaid spectra is shown in Supplementary Figure S7). **(c)** Cartoon of the G-quadruplex G4-TEMPO construct used for paramagnetic relaxation enhancement (PRE) NMR. The normalized oxidized versus reduced intensity ratios from HSQC spectra shows signal attenuation in different regions in the ZNF706 due to proximity to the TEMPO moiety. Asterisks denote unassigned resonances. Strong PRE effects are localized to the N-terminal residues, consistent with direct G-quadruplex interaction. **(d)** Size exclusion chromatography (SEC) profile of GGGGCC repeat G-quadruplex (4-repeat) showing three major species (*, **, ***) eluting at distinct volumes. **(e)** Circular dichroism spectra of SEC-resolved G-quadruplex fractions reveal structural polymorphism: * corresponds to a major hybrid, ** to parallel G-quadruplex, and *** to antiparallel topology. The titration of ZNF706 into these different isolated GGGGCC G-quadruplex topology shows ZNF706 binding to the parallel G-quadruplex structures reduces the characteristic CD signal at 264 nm, consistent with G-quadruplex destabilization. **(f)** Fluorescence-based G-quadruplex destabilization assay using a 4-repeat GGGGCC G-quadruplex labeled with 5′ FAM and 3′ BHQ1 quencher. Increasing concentrations of ZNF706 or DHX36 helicase induce dequenching of FAM fluorescence, indicating unwinding or conformational rearrangement of the G-quadruplex structure. BSA control shows no effect. Together, these results establish that ZNF706 binds directly to GGGGCC G-quadruplexes via its disordered N-terminus and destabilizes parallel G-quadruplex structures. All samples are prepared in in 20 mM NaPi, 100 mM KCl, pH 7.4. NMR experiments were done on an 800 MHz spectrometer at 4 °C, whereas SEC, CD and fluorescence measurements were done at room temperature.

While chemical shift perturbations locate interaction sites, they do not reveal spatial orientation. To gain geometric insight, we performed paramagnetic relaxation enhancement (PRE) experiments using site-specific nitroxide (TEMPO)-labeled GGGGCC or UUAGGG RNA (Figure 7c and S7, left panel). PRE effects cause distance-dependent signal broadening, allowing residue-level distance constraints. PRE signal broadening was observed exclusively for N-terminal residues, especially in the presence of 3′-TEMPO-labeled RNA. The zinc finger domain’s signals remained unaffected by RNA addition. Our results indicate that ZNF706’s N-terminal residues lie within ∼15 Å of the RNA’s 3′ end, while the zinc finger domain is more distal. Together with chemical shift perturbation data, these results support a polarized binding mode where the N-terminal region docks at the 3′-terminal G-quadruplex tetrad or flanking loops, with the ZnF domain positioned away.

To test whether the ZnF domain spatially reorganizes after G-quadruplex recognition by the N-terminus as observed in our previous study^35^, we used saturation-transfer difference NMR. Strong signals were found for N-terminal residues indicative of direct interaction. However, saturation of ZNF706’s Glu70 in the zinc finger domain also induced STD effects at specific G-quadruplex imino protons and vice versa, indicating proximity (Figure S7, right center panel). These effects were weaker than those seen at the N-terminus, suggesting transient or passive contacts between the ZnF domain and the RNA. Thus, while G-quadruplex binding is driven primarily by the SERF-like N-terminus, the ZnF domain may contribute weak or auxiliary interactions that may help position or stabilize the complex.

### ZNF706 binding destabilizes structural folding of GGGGCC repeats

ZNF706 engages G-quadruplexes via its disordered N-terminus and has been shown to modulate G-quadruplex structures^35^. Given the structural polymorphism of GGGGCC repeats, we asked whether ZNF706 exhibits topology-selective binding. We combined size exclusion chromatography (SEC), circular dichroism (CD), and fluorescence spectroscopy to characterize how ZNF706 interacts with and possibly remodels distinct G-quadruplex topologies formed by GGGGCC repeats^14^. These experiments aimed to elucidate whether structural destabilization by ZNF706 might contribute to its observed effects on translation suppression, condensate modulation, and RNA aggregate dissolution. We used DNA GGGGCC repeats for these studies that are more easily sourced and synthesized following the rational of Raguseo et al^51^.

The SEC profile of folded GGGGCC repeats revealed three distinct elution peaks denoted as *, **, and *** in Figure 7d. Peaks * and ** correspond to higher-order assemblies, likely associated with hybrid and parallel G-quadruplex structures (CD peak at ∼263 nm), while the dominant *** peak aligns with a monomeric antiparallel G-quadruplex fold (CD peak at ∼295 nm), consistent with GGGGCC’s previously reported structure^14^ (Figure 7e). The observation that a parallel G-quadruplex structure favors multimeric states is consistent with the higher known aggregation potential of this topology^48,50^. CD titration of ZNF706 against folded GGGGCC revealed a dose-dependent decrease in the ∼263 nm ellipticity, a wavelength that corresponds to the parallel G-quadruplex fold (Figure 7e, middle). This effect was observed for both 4-repeat and 10-repeat GGGGCC substrates (Figure S7, bottom right), while the signal at ∼295 nm (antiparallel) was minimally affected (Figure 7e, right). These results suggest that ZNF706 selectively destabilizes parallel G-quadruplexes, acting not as a general G-quadruplex binder but as a topology-sensitive modulator of G-quadruplex structure.

To quantify ZNF706 induced G-quadruplex destabilization, we used a fluorescence-based molecular beacon assay^52,53^ (Figure 7f, left), in which a 4-repeat GGGGCC sequence is labeled with 5′-FAM and a 3’ Black Hole Quencher 1 (3′-BHQ1). In the folded G-quadruplex state, FAM fluorescence is quenched, and G-quadruplex destabilization leads to strand separation and increased fluorescence. Titration of ZNF706 caused a rapid, sustained fluorescence increase similar to that produced by the known G-quadruplex ATP dependent helicase DHX36^27^ (Figure 7f, right). This result demonstrates that ZNF706 can efficiently destabilize G-quadruplexes though it does so without ATP hydrolysis, suggesting that upon binding, ZNF706 perturbs the conformational equilibrium of the G-quadruplex rather than functioning as an active helicase. Bovine serum albumin showed negligible fluorescence change, confirming the G-quadruplex remained stable in the absence of an active factor. Combined with the CD data, these outcomes suggest that ZNF706 recognizes features specific to parallel G-quadruplexes and disrupts them through a conformation-selective mechanism, distinguishing its mode of action from more broadly acting G-quadruplex remodelers like DHX36.

## Discussion

Our findings show that the microprotein ZNF706 functions as a regulator of *C9orf72* GGGGCC repeat RNA phase behavior and translation. Expanded GGGGCC RNAs autonomously assemble into higher-order condensates via multivalent G-quadruplex stacking, and progressive gel-to-solid transitions correlate with the progression of ALS and FTD^50^. Although we find that ZNF706 overexpression did not prevent the formation of nuclear GGGGCC RNA foci, it significantly increased the foci’s internal mobility in cells (Figure 2), shifting repeat RNA assemblies from immobile, solid-like aggregates towards a more dynamic, reversible state.

Mechanistically, we find that ZNF706 does not function as a passive scaffold. Biophysical and structural analyses indicate that ZNF706 binds to and alters parallel G-quadruplex conformations, weakening the RNA-RNA interactions that drive condensate solidification^54^ (Figures 5 and 7). This molecular fluidizer activity reduces dense internal packing and may permit exchange of RNA-binding proteins and dipeptide protein repeat products, enhancing accessibility to proteostatic clearance pathways. ZNF706 also directly represses GGGGCC-driven translation in vitro, consistent with a model in which G-quadruplex remodeling shifts repeat RNA into a translation-incompetent state and attenuates RAN translation.

As ZNF706 directly binds G-quadruplex elements and can dissolve G-quadruplex aggregates, we propose a model in which ZNF706 targets the repeat RNA within pathological condensates to inhibit its non-AUG translation and facilitate peptide clearance (Figure 8). By altering the conformation of G-quadruplex structures within these repeats, we propose that ZNF706 induces a translation-incompetent conformation or shifts the RNA into a condensate state incompatible with efficient ribosome recruitment, analogous to FMRP’s mechanism. By disassembling RNA-protein aggregates, ZNF706 may expose dipeptide repeat proteins to proteolytic pathways and serve as a safeguard against aberrant repeat RNA translation. This mechanism would limit the accumulation of neurotoxic dipeptide repeat proteins and mitigate the cellular stress caused by the repeat RNA expansions. ZNF706, by restoring internal mobility, may promote RNA-protein exchange and support ribonucleoprotein homeostasis. This microprotein appears to tune the material state of repeat RNA condensates at the intersection of structure, phase behavior, and unconventional translation. ZNF706 acts to affect the phase boundary between reversible and irreversible RNA condensation for at least one pathologically relevant RNA repeat.

**Figure 8.**
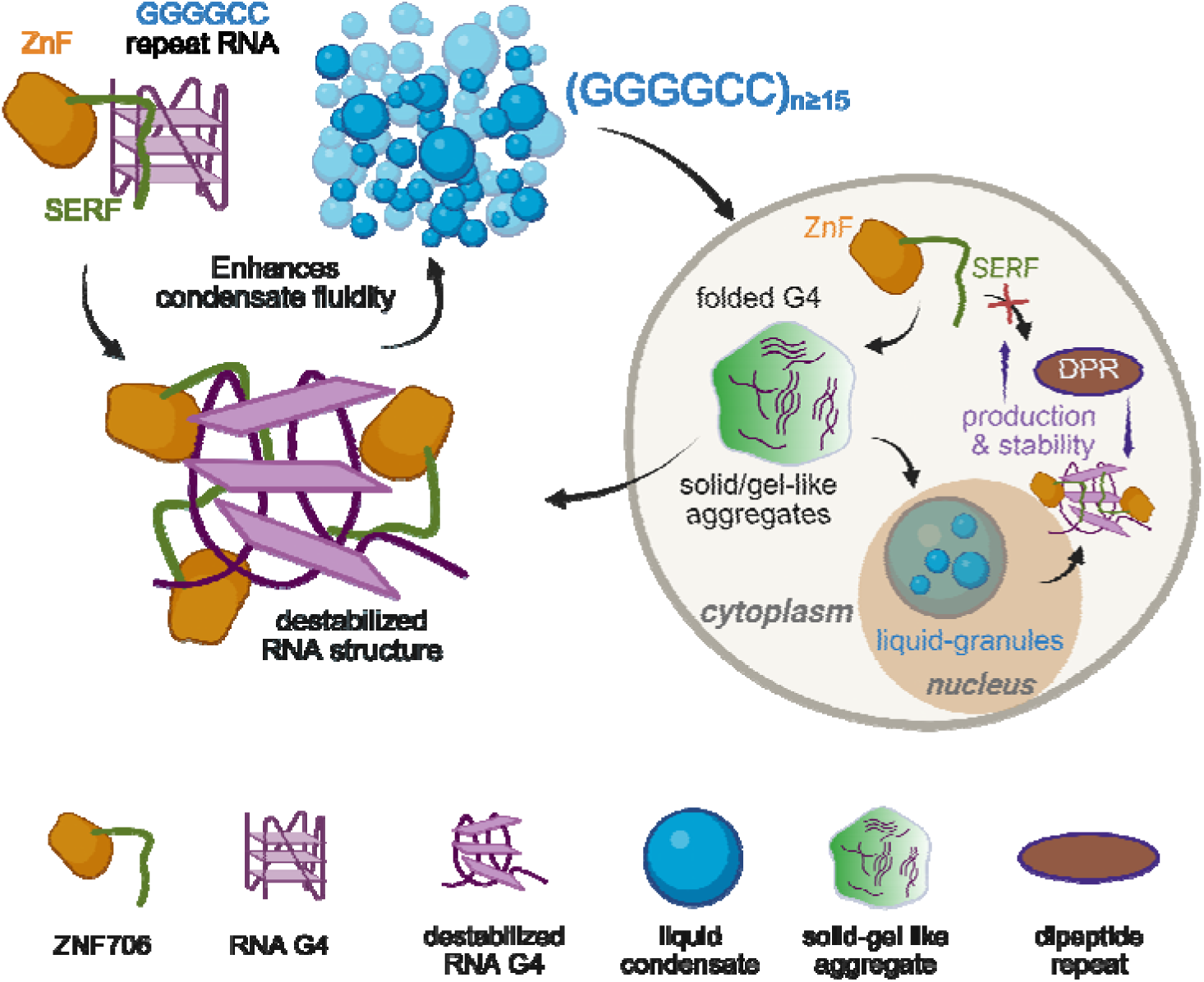
Proposed model for ZNF706 function in remodeling GGGGCC repeat RNA aggregates and in regulating phase behavior and translation. ZNF706 binds GGGGCC repeat RNA via its N-terminal SERF domain with preferential affinity for parallel G-quadruplex or G4 structures. Expanded (GGGGCC)LJ RNAs form folded G-quadruplexes that assemble into solid or gel-like aggregates in vitro and nuclear RNA foci in cells. ZNF706 converts these inert aggregates into dynamic condensates by destabilizing G-quadruplex structures, enhancing fluidity and promoting a more liquid-like phase. In cells, ZNF706 depletion increases the production and stability of poly-GA dipeptide protein repeats, suggesting ZNF706 limits RAN translation and dipeptide protein repeats turnover. The model integrates in vitro biophysical data with cellular assays, positioning ZNF706 as a G-quadruplex targeting ribo-nucleic acid particle remodeler that dissolves pathogenic RNA assemblies and safeguards cells by promoting condensate fluidity and limiting aberrant translation. Figure created with BioRender.com.

In contrast to factors that regulate repeat RNA export or translation, such as helicase DDX3X^55^ and SRSF1^56^, ZNF706 appears to shift GGGGCC repeat RNA into a less translationally active state. Several RNA-binding proteins have been implicated in modulating *C9orf72* repeat RNA metabolism, including nucleolin^14^ and Pur-α ^57^, which bind GGGGCC repeats and influence RNA toxicity, GGGGCC localization, and nuclear foci formation in disease models. However, unlike these larger RNA-binding factors for which structural information is difficult to obtain, ZNF706 represents a minimal, biophysically amenable system linking RNA structure, condensate behavior, and repeat-associated translation. ZNF706 lacks ATPase or helicase domains yet displays RNA chaperone-like activity, directly remodeling G-quadruplex rich RNA assemblies.

More broadly, length dependent G-quadruplex driven solidification and fusion arrest of repeat RNA condensates may nucleate persistent ribonucleoprotein assemblies that scaffold neurotoxic phase transitions^54^. ZNF706 functions by limiting both condensate hardening and dipeptide repeat production gain-of-function toxicities. A small, single zinc finger protein not previously linked to *C9orf72* pathology, ZNF706 highlights how structurally amenable microproteins can exert control over repeat RNA aggregation and translation, suggesting new avenues for therapeutic modulation of repeat expansion disorders.

## Methods

### Oligonucleotide preparation and labeling

Unlabeled and fluorescently labeled (Cy5 or 6-FAM) RNA or DNA oligonucleotides were obtained from Integrated DNA Technologies (IDT). The synthesized sequences included a variety of repeat sequences of defined lengths (GGGGCC 4, 6, 10, 15, 20, 32 repeats), 4-repeat UUAGGG, 4-repeat UUACCC, 4-repeat CCCCGG, and 6-repeat UGGGGU (average molecular weight ∼900 kDa). Random-length polyA (Cat.# P9403), polyU (Cat.# P9528), polyG (Cat.# P4404) and polyC (Cat.# P4903) RNA was procured from Sigma-Aldrich.

All oligonucleotides were dissolved in buffer prepared with nuclease-free water (Invitrogen, Cat.# AM9937). To ensure purity and removal of small molecule contaminants, RNAs were desalted using Amicon® Ultra 0.5□mL cutoff filters, 3kDa filters (Cat# UFC500324) for 4 or 6 GGGGCC repeats and 10□kDa (Cat# UFC9010) for 15, 20 or 32 repeats. The desalting protocol included a pre-treatment incubation with 20□mM Tris-HCl at pH□>□11 at 95□°C for 3□minutes, followed by ten rounds of buffer exchange with nuclease free water. Final RNA or DNA concentrations were determined spectrophotometrically on a Nanodrop-1000 spectrophotometer using extinction coefficients calculated with the IDT OligoAnalyzer™ tool.

To fold these repeat sequences in a way that either favored or disfavored G-quadruplex formation, samples were annealed in either potassium-containing 20□mM potassium phosphate (KPi) pH□7.4, 100□mM KCl to favor G-quadruplex formation or 20□mM sodium phosphate (NaPi), pH□7.4, 100□mM NaCl to disfavor G-quadruplex structures using a thermocycler starting at 95 °C and ramped down at 1□°C per minute. Annealed RNA and DNA sequences were stored at 4□°C for immediate use or aliquoted and frozen at -20□°C for long-term storage.

### Synthesis of long GGGGCC RNA repeats

Plasmids encoding 35-repeats of GGGGCC or 69-repeats of GGGGCC with one repeat mutated to (GGGGCA, Table S1) fused with a flanking intronic region were generously provided by Peter Todd at the University of Michigan. These plasmid inserts were recloned to pcDNA3.1(+) to remove the intronic sequence and to enable repeat GGGGCC in vitro RNA transcription. For these in vitro transcription reactions, the plasmid was digested with NdeI and AgeI restriction enzymes to isolate the T7 promoter and 0.5□μg of linearized plasmid DNA was used as a template in reactions set up with the T7 RiboMAX™ Express Large Scale RNA Production system kit (Promega, Cat.# P1320), following the manufacturer’s instructions. Post-transcription, samples were treated with DNase I at 37□°C for 15□minutes to remove residual DNA. RNA was purified using MicroSpin® G25 columns (GE Healthcare Cat.#, 27-5325-01) following manufacture instructions and folded in 20 mM KPi, 100 mM KCl, pH 7.4.

### Expression and purification of recombinant proteins and fluorophore labeling

Recombinant human ZNF706 and the ZNF706 A2C mutant, which was used for fluorescent labeling, were expressed and purified as previously described,^35^ using a pET28a-SUMO vector system in *E. coli* BL21(DE3). For NMR experiments, ^15^N- or uniformly ^15^N/^13^C-labeled ZNF706 was expressed in M9 minimal media with isotope-labeled precursors. Final purification included HisTrap affinity, SUMO protease cleavage, ion exchange (HiTrap SP), and size exclusion chromatography (Superdex 75), with buffer exchange to 40□mM HEPES (pH 7.5). Protein concentration was quantified spectroscopically (Shimadzu UV-1900) using absorbance at 280 nm and an extinction coefficient of 1490 M^-1^. cm^-1^. The 19.4 kDa fragment of the TDP-43 protein containing the RRM1-RRM2 domain (residue 102-269) was expressed and purified as described for ZNF706. The ZNF706 concentration was measured spectroscopically using a predicted extinction coefficient at 280 nm of 15720 M^-1^. cm^-1^.

For site-specific fluorescent labeling, 200□µM of the ZNF706 A2C mutant was incubated with a 10-fold molar excess of Cy5 (Cytiva, PA25031) or Alexa Fluor 488 maleimide (Invitrogen, A10254) in 20□mM Tris-HCl, 0.2□mM ZnCl_2_, 100 mM KCl, pH 8.0. The Cy5 and Alexa labelled proteins were used to perform in vitro phase separation and fluorescence polarization assays, respectively. The reaction was carried out overnight at 25□°C with 300 rpm shaking in the dark. Unreacted dye was removed via PD-10 desalting columns, and labeled protein was concentrated using 3□kDa Amicon filters (Millipore). Residual free dye was eliminated by repeated buffer exchange (8× 15□min, 7000×g) into the 20 mM NaPi, pH 7.4, 100 mM KCl buffer.

### NMR spectroscopy and PRE experiments

All NMR experiments were performed in 20□mM NaPi, 100□mM KCl, pH□7.4, with 10% D_2_O at 4□°C. 100□μM ^15^N-labeled ZNF706 was titrated with increasing concentrations of folded GGGGCC or TERRA (UUAGGG) G-quadruplex RNAs to monitor chemical shift perturbations (CSPs) via 2D ^1^H-^15^N Heteronuclear Single-Quantum Correlation (HSQC) spectra. Paramagnetic relaxation enhancement (PRE) experiments were conducted using 50□μM 2,2,6,6-tetramethylpiperidine 1-oxyl (TEMPO) labeled RNA G-quadruplexes mixed with 100□μM ZNF706 in para- and dia-magnetic conditions. PRE effects were evaluated by monitoring signal intensity changes in HSQC spectra. A 20× molar excess of sodium ascorbate was used for reduction of the paramagnetic label. Saturation transfer difference NMR experiments were carried out using 100□μM of 2-repeat UUAGGG RNA mixed with or without unlabeled ZNF706, focusing on well-resolved guanine imino protons and selected amide signals. On-resonance saturation was applied to guanine protons, and off-resonance saturation was applied at - 40□ppm, well outside the observed ^1^H spectral range to control for non-specific effects. NMR spectra were acquired on a Bruker 800□MHz spectrometer equipped with a triple resonance inverse detection TCI cryoprobes. Data was processed using Topspin 4.1.4 and analyzed in NMRFAM-Sparky v1.47.

### Biophysical measurements

#### Fluorescence polarization binding assay

Fluorescence polarization assays were conducted using Alexa Fluor 488 (AF488) labeled ZNF706 A2C mutant protein to monitor ZNF706 binding to repeat RNA. 200□nM of the labeled protein was mixed with increasing concentrations of GGGGCC repeat of varying lengths (4, 6, 10, 15, 32 and 69 repeats) folded in different conditions. For the ZNF706 binding assay using 6-repeat UGGGGU RNA, 200 nM AF488-ZNF706 was used with increasing RNA G-quadruplex concentration. All samples were prepared in nuclease-free buffer and incubated for 30□min at room temperature to allow binding equilibrium. The assay measurements were performed using a TECAN Infinite M1000 microplate reader at 25□°C with excitation/emission settings of 470/530□nm for AF488. Data were analyzed using GraphPad Prism v10.4.2 with non-linear regression fitting to a one-site binding model for apparent K_D_ determination when applicable.

#### Size-Exclusion chromatography

To isolate distinct topological species of GGGGCC repeat DNA, 20□µM of 4- or 10-repeat GGGGCC was prepared in 20□mM NaPi and 100□mM KCl pH 7.4 and incubated at room temperature for 30□min to allow proper folding. The G-quadruplex samples were then subjected to size-exclusion chromatography using a Superdex 200 Increase 10/300 GL column (Cytiva, 28-9909-44) equilibrated in the same buffer. Eluted fractions were collected and subsequently analyzed by circular dichroism (CD) spectroscopy to characterize their secondary structures.

#### Circular dichroism spectroscopy

Structural changes in GGGGCC repeat were monitored by CD spectroscopy upon titration with ZNF706. CD spectra were collected at 25□°C using a JASCO J-1500 spectropolarimeter in 20□mM NaPi and 100□mM KCl pH 7.4. 5□µM folded GGGGCC G-quadruplex was titrated with increasing concentrations of ZNF706 ranging from 1 to 5 equivalents, and spectra were recorded from 200 to 320□nm. Each spectrum represents an average of 8 scans, and baseline correction was done by subtracting buffer-only spectra. Changes in the G-quadruplex parallel and antiparallel peaks at ∼263 and 295□nm, respectively, were used to assess conformational alterations induced by ZNF706 binding.

#### Electrophoretic Mobility Shift Assay (EMSA)

Electrophoretic mobility shift assays were performed using 5□µM of 4-repeat GGGGCC or UUAGGG repeat RNA incubated with increasing concentrations of ZNF706 (2.5, 5 and 10□µM) in a binding buffer of 20□mM NaPi, 100□mM KCl, pH 7.4. In another assay, 0.5□µM of 20-repeat GGGGCC was mixed with varying ZNF706 concentrations (0.25, 0.5, 1 and 2.5 µM). Samples were incubated for ∼30 minutes at room temperature, diluted with loading dye by half, and loaded onto a 4-20% native TBE polyacrylamide gel (Invitrogen, EC6225) and electrophoresed at 130□V for 1 hour at room temperature. Gels were post-stained with SYBR Safe (ApexBio, A8743) and imaged using a Bio-Rad Gel Doc to visualize RNA-protein complexes.

#### Static Fluorescence Spectroscopy

Static fluorescence measurements were conducted using a Horiba FluoroMax 4 spectrofluorometer. A 4-repeat GGGGCC oligonucleotide labeled with a 5′ BHQ1 quencher and a 3′ 6-FAM fluorophore was synthesized by IDT and folded in 20□mM NaPi, 100□mM KCl, pH□7.4 buffer. Measurements were performed at 25□°C using 50□nM of folded G-quadruplex, excited at 495□nm with emission recorded at 517□nm. Fluorescence intensity was measured in the absence or presence of DHX36 helicase (50□nM, Europrotein cat# EP8694911), ZNF706 (100□nM and 250□nM), or BSA (100□nM, Sigma-Aldrich, A7906) to assess G-quadruplex conformational changes.

### In vitro phase separation and disaggregation assays

Phase separation experiments were performed in 16-well plates (Grace Bio-Labs, Cat.# 112358) pre-treated with 5% w/v Pluronic™ F-127 (Sigma, P2443) for 4 hours at 37 °C to prevent non-specific adhesion. Wells were rinsed three times with double-deionized water followed by working buffer of 20□mM NaPi, 100□mM KCl, pH 7.4 and air-dried. Phase separation was induced by incubating 50□µM ZNF706 with 10□ng/µL total HeLa cell RNA or synthetic 4-repeat or 15-repeat GGGGCC in working buffer that contained 10% w/v PEG8000. Other repeat RNAs that include a 4-repeat UUAGGG and a 6-repeat UGGGGU were also used for phase separation assay with ZNF706. Fluorescent imaging was carried out using 1:200 molar ratio of Cy5-labeled ZNF706 or 6-FAM labeled RNA. FRAP measurements were conducted on Cy5-labeled ZNF706 for HeLa cell RNA, 4-repeat and 15-repeat GGGGCC condensates, using a Nikon Ti2-E inverted microscope equipped with a SOLA 365 LED and an 100X oil immersion objective. Recovery kinetics were analyzed in GraphPad Prism.

For disaggregation assays, GGGGCC repeats (8, 15, 20, 35 and 69) were first thermally folded in working buffer in the presence of 10% PEG8000 and imaged to confirm aggregate formation. For short-repeat RNA, the disaggregation was initiated by adding 5- to 10-fold molar excess of ZNF706 containing 1:200 Cy5-ZNF706 to unlabeled ZNF706 and monitored over time by fluorescence microscopy. FRAP analysis of ZNF706 induced disaggregate condensates or aggregates were measured on sample containing 0.5 µM 35- or 69-repeat GGGGCC RNA and 20µM ZNF706. The effect of increasing concentration of ZNF706 varying from 10 µM to 100 µM against 0.5 µM 69-repeat RNA was also tested by incubating sample mixtures for ∼30 minutes at room temperature. For small molecule modulation, 2□µM unfolded 15R GGGGCC was incubated with 20□µM pyridostatin (Sigma-Aldrich, Cat.# SML2690) to induce aggregation. Disaggregation was subsequently triggered by adding 20□µM ZNF706. In separate experiments, 10□µM unfolded 15-repeat RNA was incubated with 20□µM TDP-43 RRM1-RRM2 domain, followed by addition of ZNF706 at 1:1 or 1:4 TDP43:ZNF706 molar ratios to assess disaggregation.

### Single particle tracking and droplet fusion by optical tweezers

Droplets were formed by mixing ZNF706 with 4- or 15-repeat GGGGCC RNA in 20□mM NaPi, 100□mM KCl pH□7.4 working buffer containing 10% w/v PEG8000. 500□nm carboxylated polystyrene beads (ThemoFisher Scientific, Cat#G500) were prepared by suspending 1 drop of bead solution as supplied by the manufacture in 1□mL in working buffer, bath-sonicated for 20□min, and centrifuged at 6000□rpm for 15□min. Five microliters of the supernatant were diluted in 995□µL of working buffer, and 1□µL of this diluted bead solution was added to the droplet mixture. An oxygen-scavenging system containing 1□mM Trolox, 2.5□mM protocatechuic acid (PCA), and 7□μg/mL protocatechuate-3,4-dioxygenase (PCD) was added to samples immediately before imaging. Droplets were introduced into a chamber made by mounting a coverslip on a glass slide using double-sided tape. The chamber was pre-washed with working buffer containing, Trolox PCA, PCD, and 10% PEG8000. After introducing the sample, the chamber was incubated upside down at room temperature for 10□min to allow droplet settling. Single particle imaging was performed using Total Internal Reflection Fluorescence or TIRF microscopy. In separate experiments, the diffusion of ZNF706 was measured within 4 or 15 repeat condensates containing ZNF706 spiked with picomolar concentration of Cy5 labelled ZNF706 for detection. The diffusion coefficients of beads or Cy5-ZNF706 were measured through single particle tracking in ImageJ following the mean squared displacement analysis using a custom MATLAB pipeline as described previously^32^.

The fusion of ZNF706 and GGGGCC 4, 6, 15 and 20 repeat droplets were done using optical tweezers (C-Trap LUMICKS). Phase separated droplets were also formed by mixing ZNF706 with random length polyRNA (polyA, polyU, polyC and polyG) or mixing ZNF706 with repeat GGGGCC prepared in 20□mM sodium phosphate pH□7.4 containing either 100□mM KCl or 100 mM NaCl.

Droplet samples were prepared in a similar way with the omission of beads and oxygen-scavenging reagents. Fusion of droplets was measured using a LUMICKS C-Trap dual-trap optical tweezer system. Two droplets were independently trapped and brought into contact until fusion was observed. Bright-field fusion events were recorded at room temperature, and fusion times were analyzed using lakeview program and fitting the data in GraphPad prism.

### Cell culture, transfection, and treatments

HEK293 cells were maintained in Dulbecco’s Modified Eagle Medium (Fisher Scientific, 11-995-073) supplemented with 10% fetal bovine serum (Sigma, F4135) and 1× Penicillin-Streptomycin-Glutamine (Fisher Scientific, 10-378-016) at 37□°C under a 5% CO_2_ atmosphere. Cells were seeded at a density of ∼ 50,000 cells per well in 24-well plates and used for dipeptide repeat proteins analysis and cycloheximide chase assays. For plasmid expression of poly-GA constructs, cells were transfected with GA3, GA35, or GA70 plasmids using Lipofectamine^TM^ LTX and PLUS^TM^ reagent (Fisher Scientific, 15338030) following the manufacturer’s instructions. Knockdown of ZNF706 was achieved by transfecting 13□nM of SMARTpool siRNA targeting ZNF706 (Horizon Discovery, M-021025-01-0005) or a non-targeting control siRNA (Horizon Discovery, D-001206-13-20) using Lipofectamine RNAiMAX (Thermo Fisher, 13778150). For detection of poly-GA dipeptide repeat proteins, anti-FLAG antibody (for GA3 constructs, Sigma-Aldrich, Cat.# F1804) or poly-GA-specific antibodies (for GA35 and GA70, Proteintech, Cat.# 24492-1-AP) were used. For ZNF706 overexpression assays, cells were transduced with a lentiviral vector expressing ZNF706 for 24 hours following GA70 transfection. Cells were then lysed at the indicated time points in 1× reducing SDS sample buffer (Bio-Rad), boiled at 95 °C for 5 min, and processed for western blotting to assess dipeptide repeat protein levels.

For cycloheximide chase assays a ZNF706 knockdown was performed by transfecting 13□nM of siZNF706 or non-targeting control siRNA using Lipofectamine RNAiMAX for 48□h. After siRNA transfection, cells were transfected with GA70 plasmid using Lipofectamine™ LTX and PLUS™ reagent and incubated for an additional 48□h to allow dipeptide repeat proteins expression. Next, cycloheximide (Sigma-Aldrich, C7698) was added to the culture medium at a final concentration of 50□µg/mL. Cells were harvested at 0, 8, 24 and 48□h following cyclohexamide addition to assess poly-GA dipeptide repeat protein stability in ZNF706-depleted versus control conditions.

Horseradish peroxidase (HRP)-conjugated Goat anti-Rabbit IgG secondary antibodies (Thermo Scientific, Cat.# 31460) were used for western blot detection, developed using the SuperSignal™ West Pico PLUS Chemiluminescent Substrate (Thermo Scientific, Cat.# 34580) were used for chemiluminescent detection in western blotting. Each experiment was performed in three biological replicates unless stated otherwise.

### In vitro translation assay

In vitro translation assay was performed using rabbit reticulocyte lysate (RRL; Promega, L4960) as previously described^34^. Briefly, 30 fmol of reporter mRNA was added to 10 µl RRL supplemented with 0.5 mM magnesium acetate, 100 mM KCl, and 1 U murine RNase inhibitor (NEB). Reactions were incubated with the indicated concentrations of ZNF706 at 30 °C for 30 min in a 96-well format. Luciferase activity was measured using NanoGlo substrate (Promega, N1110) on a Tecan M1000 plate reader. Translation products were further analyzed by anti-FLAG immunoblotting following sample denaturation by heating at 95°C for ∼5 minutes in SDS sample buffer.

### Immunofluorescence and live-cell imaging

A monoclonal U2OS cell line stably expressing Tet3G and YFP-MS2 reporter constructs (gift from Dr. Ankur Jain, Whitehead Institute) was used for visualizing nuclear RNA foci following a minor adaptation of the methods described previously^5^. Briefly, cells were seeded in 8-well chambered cover glass slides (Ibidi, 80806) and transduced with a lentiviral vector expressing MS2-tagged 34× GGGGCC repeats or 23×UUAGGG repeats under a doxycycline-inducible promoter. Cells were seeded at ∼50,000 in 200 µL culture media overnight, followed by 40 µL of lenti titer treatment. Transductions were performed in the presence of 8□μg/mL polybrene for 24□h. After 24□h of lentiviral transduction, cells were transiently transfected with pcDNA3.1 ZNF706-mCherry plasmid using Lipofectamine^TM^ LTX and PLUS^TM^ reagent according to the manufacturer’s protocol as described above. Following an additional 48□h, doxycycline was added (2□μg/mL final concentration) for 48□h to induce repeat RNA expression. Live-cell imaging was performed using a Leica SP8 confocal microscope using a 63× oil-immersion objective. For nuclear staining, cells were incubated with NucBlue™ Live ReadyProbes™ reagent (Invitrogen, R37605) according to the manufacturer’s instructions. Fluorescence signals were collected using sequential excitation and emission filters suitable for mCherry, YFP, and DAPI. Image acquisition and processing were performed using Leica LAS X and Fiji ImageJ software. Cell passage numbers between 7 and 15 were used for all imaging experiments, and ∼50,000 cells were seeded per well to maintain consistent confluency. Imaging of YFP-positive nuclear RNA foci was conducted under live conditions at 37□°C using an environmental chamber.

## Acknowledgements

This work was supported by the Howard Hughes Medical Institute (HHMI) where J.C.A.B. is an Investigator. This work was also supported by the National Institute of Health grant R35 GM122506 to U.J. Salary was paid from NIH for J.B. The salary for B.S. was provided by the Howard Hughes Medical Institute. The funders had no role in study design, data collection and analysis, decision to publish, or preparation of the manuscript. Plasmids containing GGGGCC repeat sequences were generously provided by Dr. Peter Todd (University of Michigan). We thank Ankur Jain (Whitehead Institute) for providing the monoclonal U2OS cell line used for studies of repeat RNA aggregation.

