## Supplemental information Figures and Tables for "Microprotein Regulates G-quadruplex Driven RNA Aggregation"

**Supporting Figures**

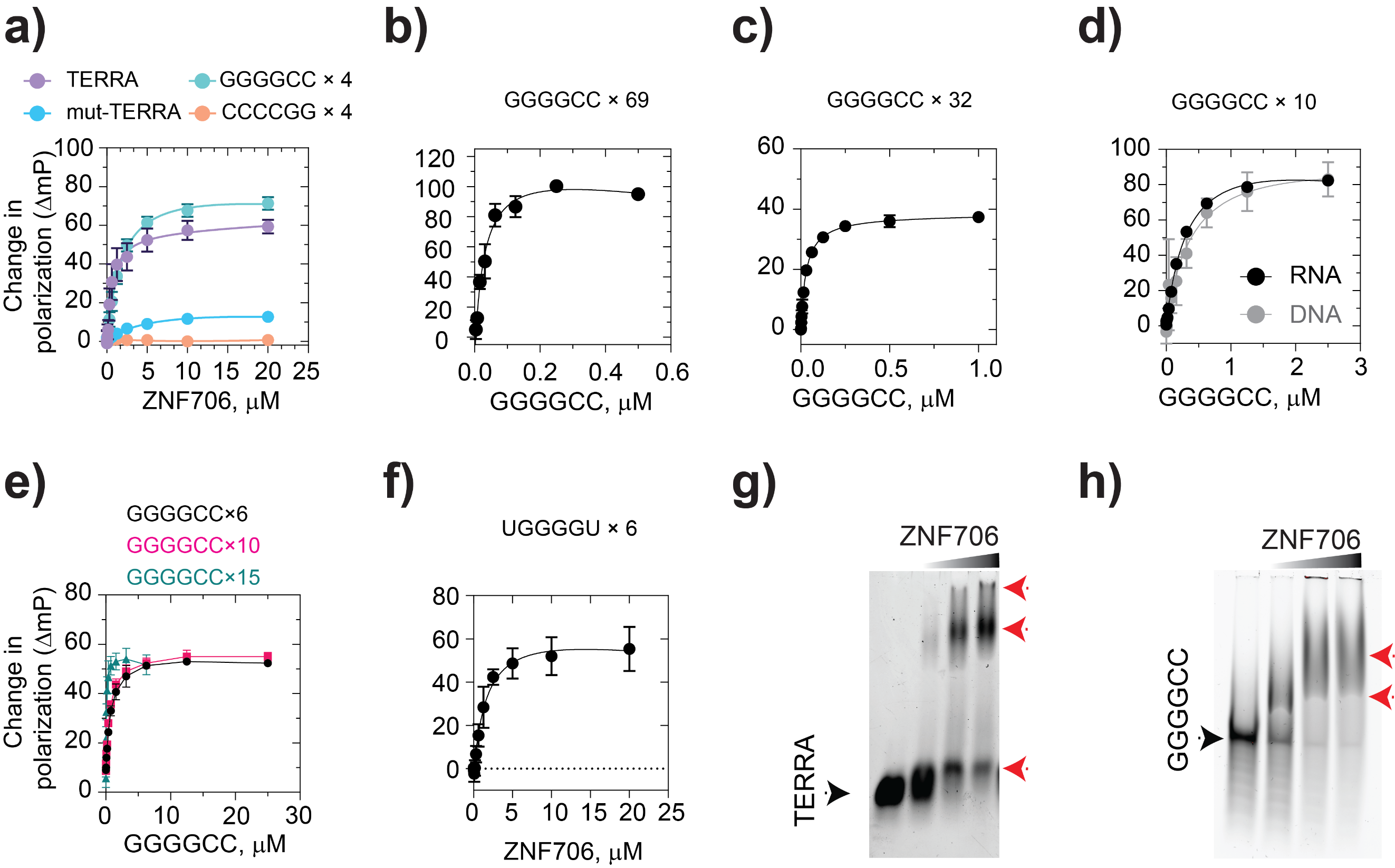

**Figure S1**. Binding of ZNF706 to GGGGCC repeats becomes stronger as the repeat number increases. **(a-f)** Fluorescence polarization binding curves of ZNF706 for different repeat-length RNA and DNA G4 quadruplexes and their corresponding non-G4 mutated sequences. RNA or DNA sequences shown in panels **(a)** have 4-repeat GGGGCC sequences, and a 6-repeat UGGGGU sequence was used in panel **(f).** Data shown as mean ± SD (n ≥ 3); affinities are in the sub-micromolar to nanomolar range. The binding affinity (Kd) values for Figure S1a-f are reported in Table 1. **(g,h)** Native EMSA of 4-repeat TERRA **(g)** or GGGGCC **(h)** RNA G-quadruplex with increasing ZNF706 concentration. Black arrowhead marks the free RNA band, red arrowheads mark ZNF706 bound RNA complexes. The gel demonstrates concentration-dependent complex formation, corroborating fluorescence polarization results. All FP and EMSA samples are prepared in 20 mM NaPi, 100 mM KCl, pH 7.4, and incubated ~30 minutes before measurement.

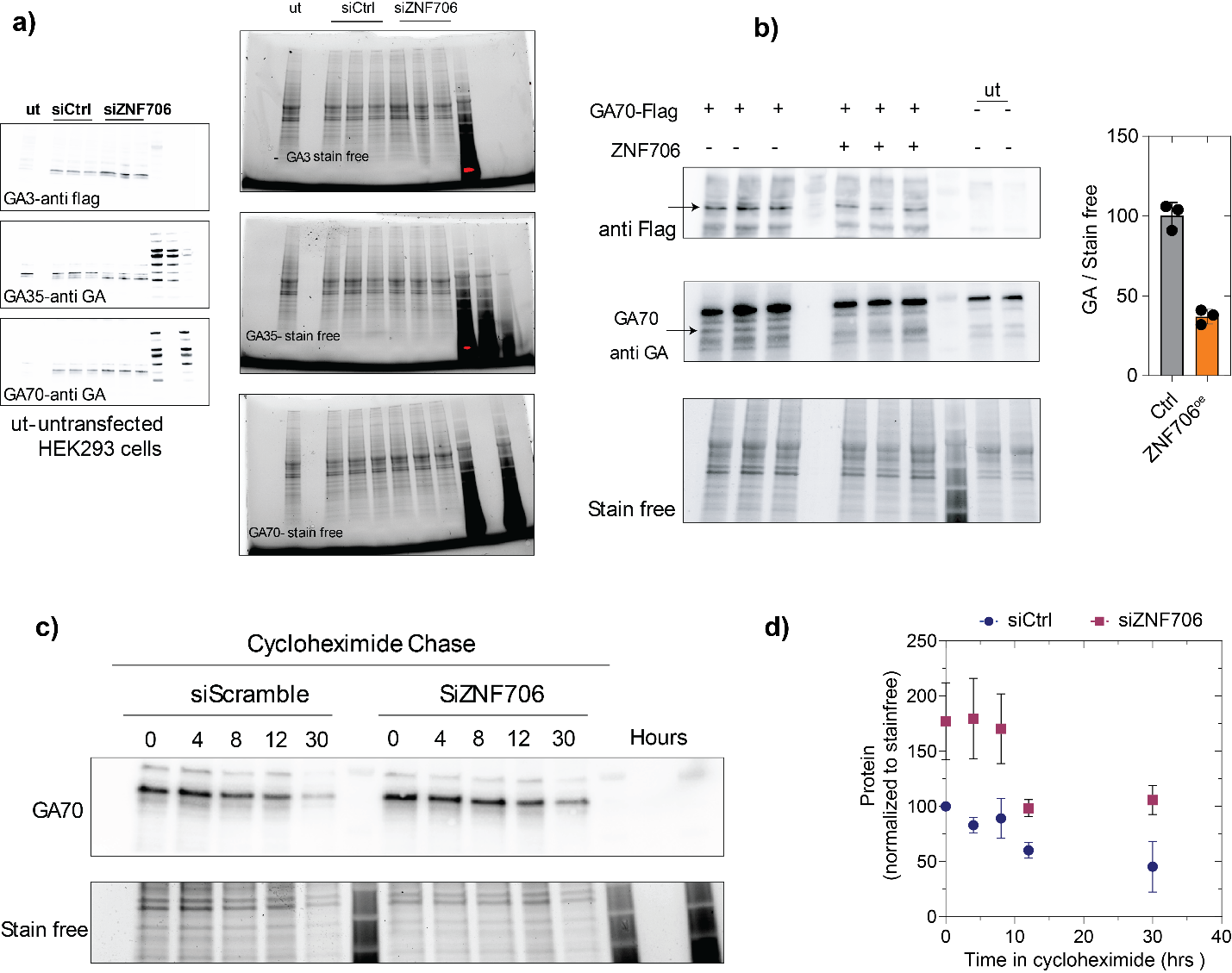

**Figure S2.** **ZNF706 suppresses repeat-RNA mediated dipeptide production.** **(a)** Immunoblots showing levels of poly-GA dipeptide repeat proteins GA3, GA35, and GA70 in HEK293 cells following control knockdown (siCtrl) or ZNF706 depletion (siZNF706). Knockdown of ZNF706 increases the accumulation of repeat-associated poly-GA species. Stain-free gel images on the right show the total protein levels in HEK293 cells expressing poly-GA constructs producing GA3, GA35, or GA70 dipeptide repeat proteins. Total protein staining was used for normalization in the quantification of poly-GA levels following control knockdown (siCtrl) or ZNF706 depletion (siZNF706), corresponding to the analyses shown in Figure 3a. **(b)** Representative immunoblots of GA70-Flag expressed in HEK293 cells in the presence or absence of ZNF706 overexpression, probed with anti-FLAG or anti-GA antibodies. Stain-free total protein staining is shown as a loading control. Quantification of GA levels normalized to stain-free signal demonstrates reduced GA abundance upon ZNF706 expression as shown in Figure 3b. **(c)** Cycloheximide chase analysis of GA70 turnover in control or ZNF706-depleted cells. Representative immunoblots show the decay of GA70 following translation inhibition over the indicated time points, with histone H3 as a loading control. **(d)** Quantification of GA70 protein levels during cycloheximide chase normalized to total protein in stain free gel. ZNF706 depletion stabilizes GA70, indicating reduced turnover. Data represent mean ± s.d., n=4.

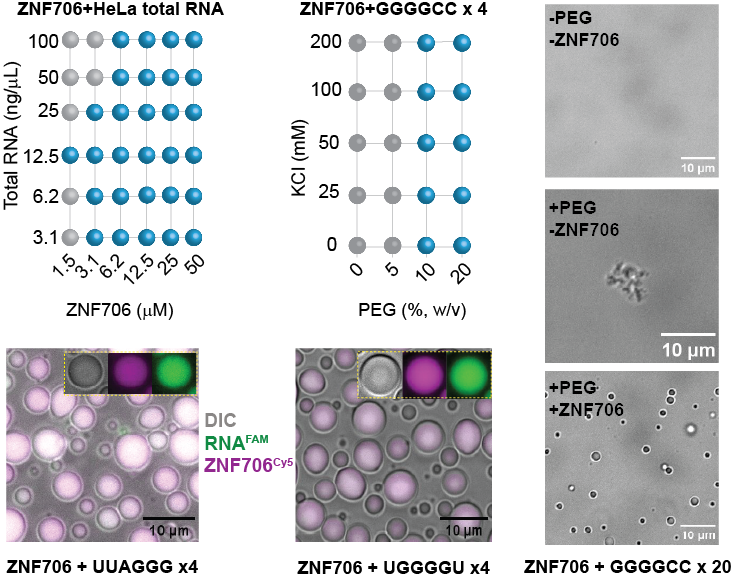

**Figure S3.** **Phase separation behavior of ZNF706 with total RNA and G4-forming GGGGCC repeats.** (Top left) Phase diagram showing that ZNF706 forms condensates with HeLa total RNA across a wide concentration range (blue circles indicate visible droplets). Condensation is robust at RNA concentrations >6.2 ng/μL and ZNF706 ≥ 3 μM. (Top middle) Phase behavior of ZNF706 with synthetic 4-repeat GGGGCC RNA as a function of PEG-8000 and KCl concentration. Droplet formation is promoted by molecular crowding (≥10% PEG w/v) and is salt-independent, consistent with G4-driven assembly. (Bottom) Confocal images of co-phase-separated droplets formed by 50 µM ZNF706 with equimolar G4-forming UUAGGG or control UGGGGU RNAs containing 1/200^th^ fraction of labeled with 5,6-FAM in green and ZNF706-Cy5 in magenta relative to unlabeled protein or RNA. Both RNAs colocalize with ZNF706, forming enriched droplets. (Top right) DIC microscopy confirms that ZNF706 does not undergo phase separation in dilute buffer (–PEG, –ZNF706), but 20-repeat GGGGCC RNA alone can form static aggregates in PEG (+PEG, –ZNF706). Upon addition of ZNF706, these aggregates convert into dynamic, spherical condensates (+PEG, +ZNF706).

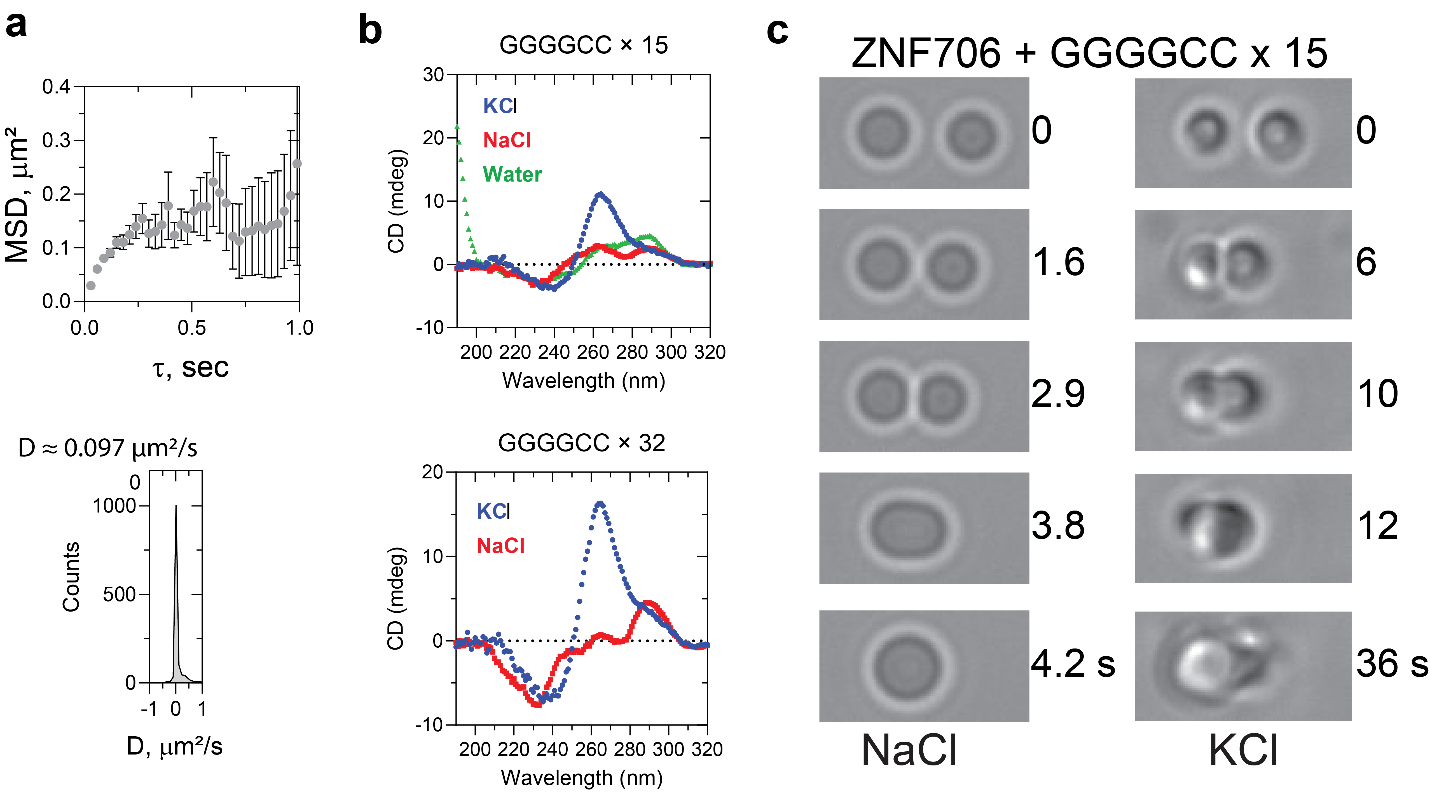

**Figure S4.** **Physical properties of ZNF706-GGGGCC condensates probed by single-molecule diffusion and optical trapping**. **(a)** Mean squared displacement (MSD) of individual ZNF706 molecules within condensates formed with 4-repeat GGGGCC RNA indicates diffusive dynamics. The apparent diffusion coefficient (D ≈ 0.097 μm²/s) was extracted from the MSD fit (middle), consistent with liquid-like behavior. **(b)** Secondary structure assessment of 15- and 32-repeat GGGGCC repeats folded in different salt or no salt conditions recorded at room temperature as indicated. **(c)** Optical tweezer fusion assay assessing material properties of droplets formed by ZNF706 with 15-repeat GGGGCC RNA under different ionic conditions. In NaCl buffer (left column), where G4 folding is disfavored, droplets readily fuse within ~4 s, indicative of low viscosity and fluidity. In contrast, under KCl conditions (right column), where G4 structures are stabilized, fusion is delayed (> 36 s), suggesting increased gel-like behavior. These results support that ZNF706 modulates GGGGCC RNA condensation and dynamics in a G4-dependent manner.

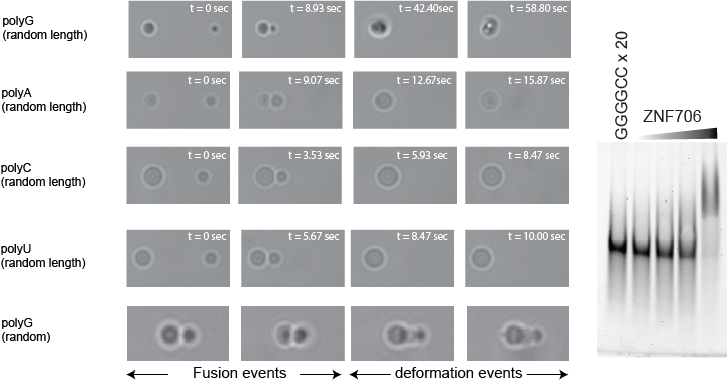

**Figure S5.** **ZNF706 interaction modulates biophysical properties of RNA condensates in a sequence-dependent manner.** (Left) Time-lapse DIC microscopy of ZNF706-driven condensates formed with various homopolymeric RNAs (random lengths of polyG, polyA, polyC, polyU, average weight ~900 kDa). Fusion and deformation dynamics were monitored following contact between droplets. PolyG exhibited markedly delayed fusion (~ > 60 s) and restricted droplet deformation (bottom row), in contrast to faster fusion events observed for polyA, polyC, and polyU (within ~10 s), indicating sequence-specific material properties. (Right) Native gel electrophoresis showing mobility shift of 20-repeat GGGGCC RNA (0.5 µM) upon titration with increasing ZNF706 concentrations (0.25, 0.5, 1 and 2.5 µM). The progressive retardation of RNA bands at high ZNF706 concentration suggests stable complex formation between ZNF706 and structured G4 RNA.

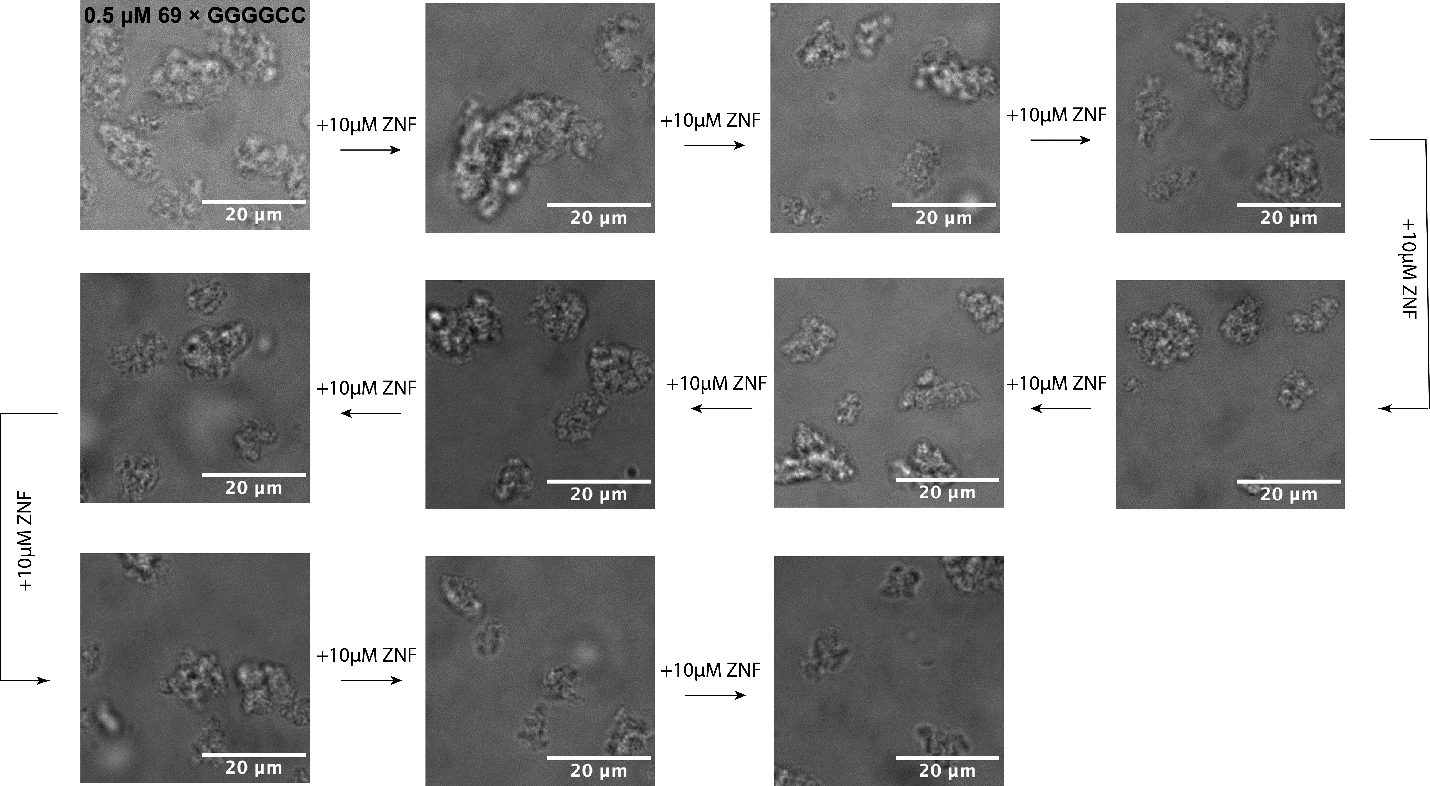

**Figure S6.** **Differential interference contrast microscopy images showing the effect of ZNF706 on solubilizing 69 × GGGGCC RNA aggregates**. 69× GGGGCC RNA (0.5 µM) aggregates were prepared in 20 mM NaPi, 100 mM KCl, pH 7.4 buffer containing 10% (w/v) PEG8000. ZNF706 was mixed to the RNA aggregates at concentrations ranging from 10 to 100 µM) and solubilization effect was monitored at room temperature.

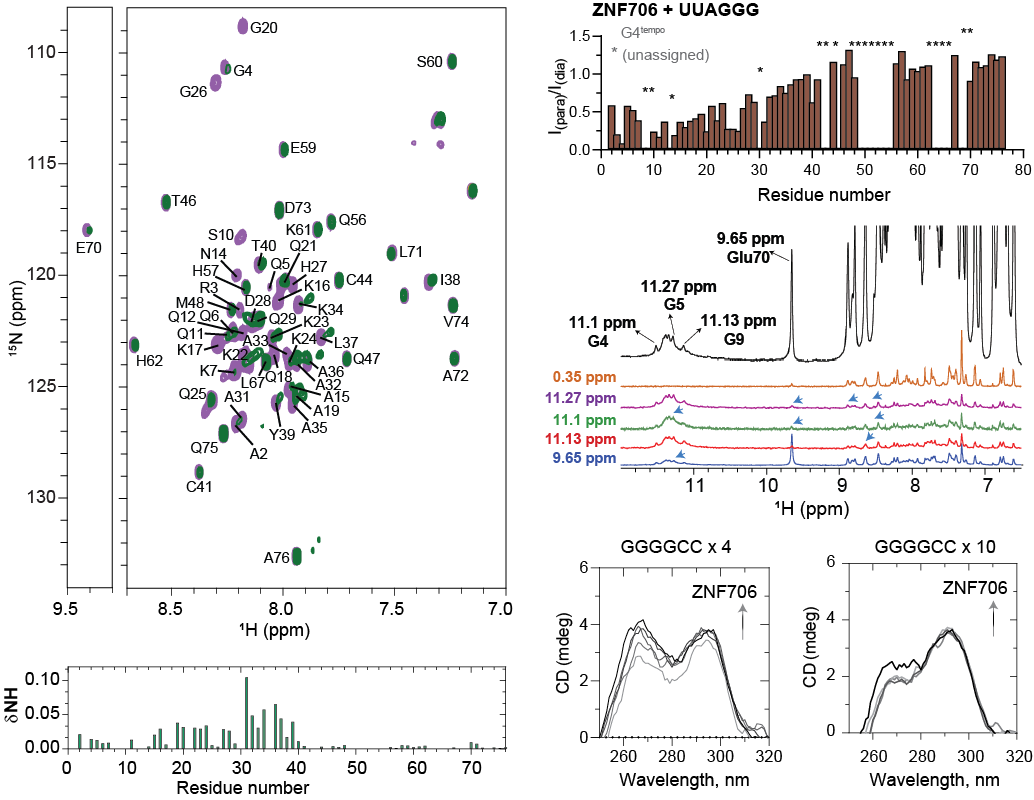

**Figure S7.** **ZNF706 selectively interacts with G4 RNA via its N-terminal domain and perturbs G4 conformation.** (Left) 1H-15N HSQC spectra of uniformly labeled ZNF706 (100 μM) overlaid in the absence (purple) and presence (green) of 4-repeat GGGGCC RNA (50 μM). Widespread chemical shift perturbations (CSPs) are observed predominantly in the disordered N-terminal residues, as quantified in the bottom histogram. (Top right) Paramagnetic relaxation enhancement (PRE) using TEMPO-labeled UUAGGG G4 RNA reveals residue-specific broadening of ZNF706 peaks under paramagnetic (I_para_) versus diamagnetic (I_dia_) conditions, showing strong PRE effects in the N-terminal region, indicating spatial proximity to the RNA’s 3′-TEMPO site. (Middle right) Saturation Transfer Difference NMR spectra probing RNA-protein interface. Selective saturation of individual UUAGGG guanine imino protons (11.27, 11.13, and 11.1 ppm) or Glu70 (ZnF domain) of ZNF706 reveals bidirectional transfer, confirming direct contact between these moieties. (Bottom) Circular dichroism spectra of 4-repeat and 10-repeat GGGGCC RNA with increasing ZNF706 concentrations. The characteristic parallel G4 peak at ~263 nm diminishes upon ZNF706 addition, suggesting partial disruption or conformational remodeling of G4 structure. All experiments were performed in 20 mM NaPi, 100 mM KCl, pH 7.4.

**Supplementary videos**

SV1. Optical tweezer based observation of fusion events in ZNF706 - 15×GGGGCC RNA condensates in KCl solution (20 mM KPi, 100 mM KCl, pH 7.4).

SV2. Optical tweezer based observation of fusion events in ZNF706 - 15×GGGGCC RNA condensates in NaCl solution (20 mM NaPi, 100 mM NaCl, pH 7.4).

SV3. Optical tweezer based observation of fusion events in ZNF706 - 4×GGGGCC RNA condensates in KCl solution (20 mM KPi, 100 mM KCl, pH 7.4).

SV4. Optical tweezer based observation of fusion events in ZNF706 - 4×CCCCGG RNA condensates in KCl solution (20 mM KPi, 100 mM KCl, pH 7.4).

SV5. Optical tweezer based observation of fusion events in ZNF706 - polyA RNA condensates in KCl solution (20 mM KPi, 100 mM KCl, pH 7.4).

SV6. Optical tweezer based observation of fusion events in ZNF706 - polyG RNA condensates in KCl solution (20 mM KPi, 100 mM KCl, pH 7.4).

SV7. Optical tweezer based observation of fusion events in ZNF706 - polyU RNA condensates in KCl solution (20 mM KPi, 100 mM KCl, pH 7.4).

SV8. Optical tweezer based observation of fusion events in ZNF706 - 20×GGGCC RNA condensates in KCl solution (20 mM KPi, 100 mM KCl, pH 7.4).

SV9. Optical tweezer based deformation of fused ZNF706 - 20×GGGCC RNA condensates by applying pull-force in KCl solution (20 mM KPi, 100 mM KCl, pH 7.4).

**Table S1.** *In vitro* synthesized GGGGCC repeat RNA sequences

| **69 × GGGGCC^¶^** | *GTGTGTGTTTTTGTTTTTCCCACCCTCTCTCCCCACTACTTGCTCTCACAGTACTCGCTGAGGGTGAACAAGAAAAGACCTGATAAAGATTAACCAGAAGAAAACAAGGAGGGAAACAACCGCAGCCTGTAGCAAGCTCTGGAACTCAGGAGTCGCGCGCTAgCGGCC* GGGGCCGGGGCCGGGGCCGGGGCCGGGGCCGGGGCCGGGGCCGGGGCCGGGGCCGGGGCCGGGGCCGGGGCCGGGGC**A**GGGGCCGGGGCCGGGGCCGGGGCCGGGGCCGGGGCCGGGGCCGGGGCCGGGGCCGGGGCCGGGGCCGGGGCCGGGGCCGGGGCCGGGGCCGGGGCCGGGGCCGGGGCCGGGGCCGGGGCCGGGGCCGGGGCCGGGGCCGGGGCCGGGGCCGGGGCCGGGGCCGGGGCCGGGGCCGGGGCCGGGGCCGGGGCCGGGGCCGGGGCCGGGGCCGGGGCCGGGGCCGGGGCCGGGGCCGGGGCCGGGGCCGGGGCCGGGGCCGGGGCCGGGGCCGGGGCCGGGGCCGGGGCCGGGGCCGGGGCCGGGGCCGGGGCCGGGGCCGGGGCCGGGGCCGGGGCCGGGGCC |
| --- | --- |
| **35 × GGGGCC^¶^** | *GTGTGTGTTTTTGTTTTTCCCACCCTCTCTCCCCACTACTTGCTCTCACAGTACTCGCTGAGGGTGAACAAGAAAAGACCTGATAAAGATTAACCAGAAGAAAACAAGGAGGGAAACAACCGCAGCCTGTAGCAAGCTCTGGAACTCAGGAGTCGCGCGCTAgCGGCC* GGGGCCGGGGCCGGGGCCGGGGCCGGGGCCGGGGCCGGGGCCGGGGCCGGGGCCGGGGCCGGGGCCGGGGCCGGGGCAGGGGCCGGGGCCGGGGCCGGGGCCGGGGCCGGGGCCGGGGCCGGGGCCGGGGCCGGGGCCGGGGCCGGGGCCGGGGCCGGGGCCGGGGCCGGGGCCGGGGCCGGGGCCGGGGCCGGGGCCGGGGCCGGGGCC |
| **69 × GGGGCC** | TATGCACGATCATGTAGTTGCGAAATTC*TAATACGACTCACTATA*GTC GGCCGGGGCCGGGGCCGGGGCCGGGGCCGGGGCCGGGGCCGGGGCCGGGGCCGGGGCCGGGGCCGGGGCCGGGGCCGGGGC**A**GGGGCCGGGGCCGGGGCCGGGGCCGGGGCCGGGGCCGGGGCCGGGGCCGGGGCCGGGGCCGGGGCCGGGGCCGGGGCCGGGGCCGGGGCCGGGGCCGGGGCCGGGGCCGGGGCCGGGGCCGGGGCCGGGGCCGGGGCCGGGGCCGGGGCCGGGGCCGGGGCCGGGGCCGGGGCCGGGGCCGGGGCCGGGGCCGGGGCCGGGGCCGGGGCCGGGGCCGGGGCCGGGGCCGGGGCCGGGGCCGGGGCCGGGGCCGGGGCCGGGGCCGGGGCCGGGGCCGGGGCCGGGGCCGGGGCCGGGGCCGGGGCCGGGGCCGGGGCCGGGGCCGGGGCCGGGGCCGGGGCCGGTCGTGGAAGGGTGGGCGCGCCCA |
| **35 × GGGGCC** | TATGCACGATCATGTAGTTGCGAAATTC*TAATACGACTCACTATA*GTC GGGGCCGGGGCCGGGGCCGGGGCCGGGGCCGGGGCCGGGGCCGGGGCCGGGGCCGGGGCCGGGGCCGGGGCCGGGGCAGGGGCCGGGGCCGGGGCCGGGGCCGGGGCCGGGGCCGGGGCCGGGGCCGGGGCCGGGGCCGGGGCCGGGGCCGGGGCCGGGGCCGGGGCCGGGGCCGGGGCCGGGGCCGGGGCCGGGGCCGGGGCCGGGGCCGGTCGTGGAAGGGTGGGCGCGCCCA |
| ^¶with an intronic sequence; bold-underlined: mutated G-A nucleotide^ | |
